# PRIZM: Combining Low-N Data and Zero-shot Models to Design Enhanced Protein Variants

**DOI:** 10.64898/2026.02.08.704628

**Authors:** David Harding-Larsen, Brianna M. Lax, Martina Escorial García, Catarina Mendonça, Felipe Mejia-Otalvaro, Ditte Hededam Welner, Stanislav Mazurenko

## Abstract

Machine learning has repeatedly shown the ability to accelerate protein engineering, but many approaches demand large amounts of robust, high-quality training data as well as substantial computational expertise. While large pre-trained models can function as zero-shot proxies for predicting variant effects, selecting the best model for a given protein property is often non-trivial. Here, we introduce Protein Ranking using Informed Zero-shot Modelling (PRIZM), a two-phase workflow that first uses a high-quality low-N dataset to identify the most suitable pre-trained zero-shot model for a target protein property and then applies that model to rank and prioritize an in silico variant library for experimental testing. Across diverse benchmark datasets spanning multiple protein properties, PRIZM reliably separated low- from high-performing models using datasets of ∼20 labelled variants. We further demonstrate PRIZM in enzyme engineering case studies targeting sucrose synthase thermostability and glycosyltransferase activity, where PRIZM-guided selection identified improved variants, including gains of ∼3°C in apparent melting temperature and ∼20% higher relative activity. PRIZM provides an accessible, data-efficient route to leverage foundation models for protein design while requiring minimal experimental data.

## Introduction

Protein engineering encompasses the design and optimization of proteins with improved or novel functions by altering their amino acid sequences. Two predominant strategies guide this process: rational design, which relies on structural and mechanistic insight, and directed evolution, which utilizes iterative cycles of mutation and selection to explore sequence space empirically (Packer and Liu 2015, Arnold 2018, Lutz and Iamurri 2018). In recent years, machine learning (ML) has emerged as a complementary approach, enabling data-driven modelling of protein sequence-function relationships (Yang, Wu, and Arnold 2019, Mazurenko, Prokop, and Damborsky 2020, Markus *et al*. 2023). ML offers the potential to generalize beyond traditional domain knowledge while still leveraging available experimental and structural data.

Pioneered by Nobel laureate Francis Arnold, ML-assisted directed evolution (MLDE) has shown impressive potential for efficiently traversing the fitness landscape of proteins, both reducing experimental demand and resulting in enhanced functionality (Yang, Wu, and Arnold 2019). In MLDE, a supervised ML-based model is trained on an initial set of labelled variants and then used to nominate new variants for experimental characterization (Biswas *et al*. 2021, Wittmann, Yue, and Arnold 2021, Hsu, Nisonoff *et al*. 2022, Zhou *et al*. 2024, Jiang *et al*. 2025, Teufel *et al*. 2025). The resulting data are iteratively fed back into the model, progressively improving its predictions. Recent MLDE methods leverage embeddings from large protein foundation models, ML-based tools trained on millions of natural sequences and structures using unsupervised objectives (Harding-Larsen, Funk *et al*. 2024). This enables the models to generalize beyond the limited size of experimental datasets (Harding-Larsen, Funk *et al*. 2024). In particular, the recently published EVOLVEpro workflow by Jiang et al. shows promise, drastically improving the hit rates of high-activity candidates using fewer than ten training rounds with only 16 mutants (Jiang *et al*. 2025).

In parallel to producing useful embeddings, large protein foundation models are used in a different mode. Because these models learn the constraints that govern natural protein families, they can provide meaningful estimates of how individual mutations affect protein properties even without an initial set of labelled variants, unlike their supervised counterparts. These models can thus be used in a zero-shot setting, where their internal scores serve as proxies for mutational effects without any task-specific training data. An example is the work by Hie et al., where the authors developed a workflow that nominates new candidates by selecting variants predicted to outperform the wildtype (WT) protein according to multiple zero-shot models (Hie *et al*. 2024).

While both supervised learning and zero-shot modelling have their benefits, both methodologies also have their own set of challenges – especially for non-experts. For supervised learning using small datasets, the most significant challenge is the susceptibility to overfitting. A robust and reliable train-test split is effectively impossible at the low-N limit, as neither subset is large enough to be statistically representative of the overall experimental dataset. To mitigate the risk, many state-of-the-art tools, such as the EVOLVEpro (Jiang *et al*. 2025), emphasize the creation of new, unbiased datasets to train robust and accurate models. However, this approach misses the opportunity to learn from existing experimental datasets, which ultimately increases experimental load and forgoes previously published data. Moreover, since supervised models are inherently tailored to specific engineering tasks, any transfer to different proteins or properties often requires model re-designing, retraining, and optimisation (Biswas *et al*. 2021), creating the need for a machine learning expert in the team. For example, how to select the most suitable pre-trained protein embedding is still an open question (Harding-Larsen, Funk *et al*. 2024). While benchmarking suites exist (e.g., TAPE (Rao *et al*. 2019)), many application papers still perform only *ad hoc* comparisons across a small number of models, and broad, systematic evaluations across the rapidly expanding set of models remain uncommon. In summary, this strong emphasis on careful model design and setup can make supervised MLDE daunting, if not downright impossible to implement for non-experts.

While zero-shot modelling appears easier to implement, it introduces its own challenges. Many models are available for zero-shot predictions, and this abundance can be a double-edged sword that makes choosing the most suitable model for a given engineering task a non-trivial challenge. Several studies have benchmarked zero-shot models against collections of deep mutational scan (DMS) datasets (Meier *et al*. 2021, Wittmann, Yue, and Arnold 2021, Notin *et al*. 2023, Li *et al*. 2024), with ProteinGym by Notin et al. being the most comprehensive to date (Notin *et al*. 2023). However, these benchmarks typically report global averages across many proteins, which do not necessarily reflect performance on the specific protein or property of interest (Hsu, Verkuil *et al*. 2022, Notin *et al*. 2023). When evaluated on subsets of tasks or proteins, the ranking of the models changes dramatically, implying that the best models may be task specific. However, zero-shot settings, by definition, do not incorporate any prior experimental data. This prevents models from adapting to specific systems or engineering objectives. As a result, model performance can vary between targets, and the best-performing model for one protein or protein property may be suboptimal for another.

Overall, these challenges highlight that neither supervised nor zero-shot approaches are straightforward for non-experts to apply in practice. Supervised methods demand careful model design and tuning, while zero-shot models offer no principled way to select the right predictor or adapt to the system at hand. This creates a need for an approach that combines the generality of foundation models with task-specific experimental insight, while avoiding specialised model setup. To address this need, we developed Protein Ranking using Informed Zero-shot Modelling (PRIZM). This novel ML approach combines the general domain knowledge of the large foundation models with the specialized insights obtained from low-N datasets to engineer enhanced protein variants. PRIZM consists of two phases: (1) Model Ranking and (2) Variant Selection (Figure 1). In the first phase, the protein information – be it sequence, structure, or evolutionary patterns – is processed by a collection of pre-trained models (Table S1). The resulting zero-shot scores are then compared to an experimental low-N dataset to identify the model that best describes the targeted protein property. In the second phase, this best model is utilized to process and rank an *in silico* library, which serves as the basis for identifying new variants for experimental characterization.

**Figure 1.**
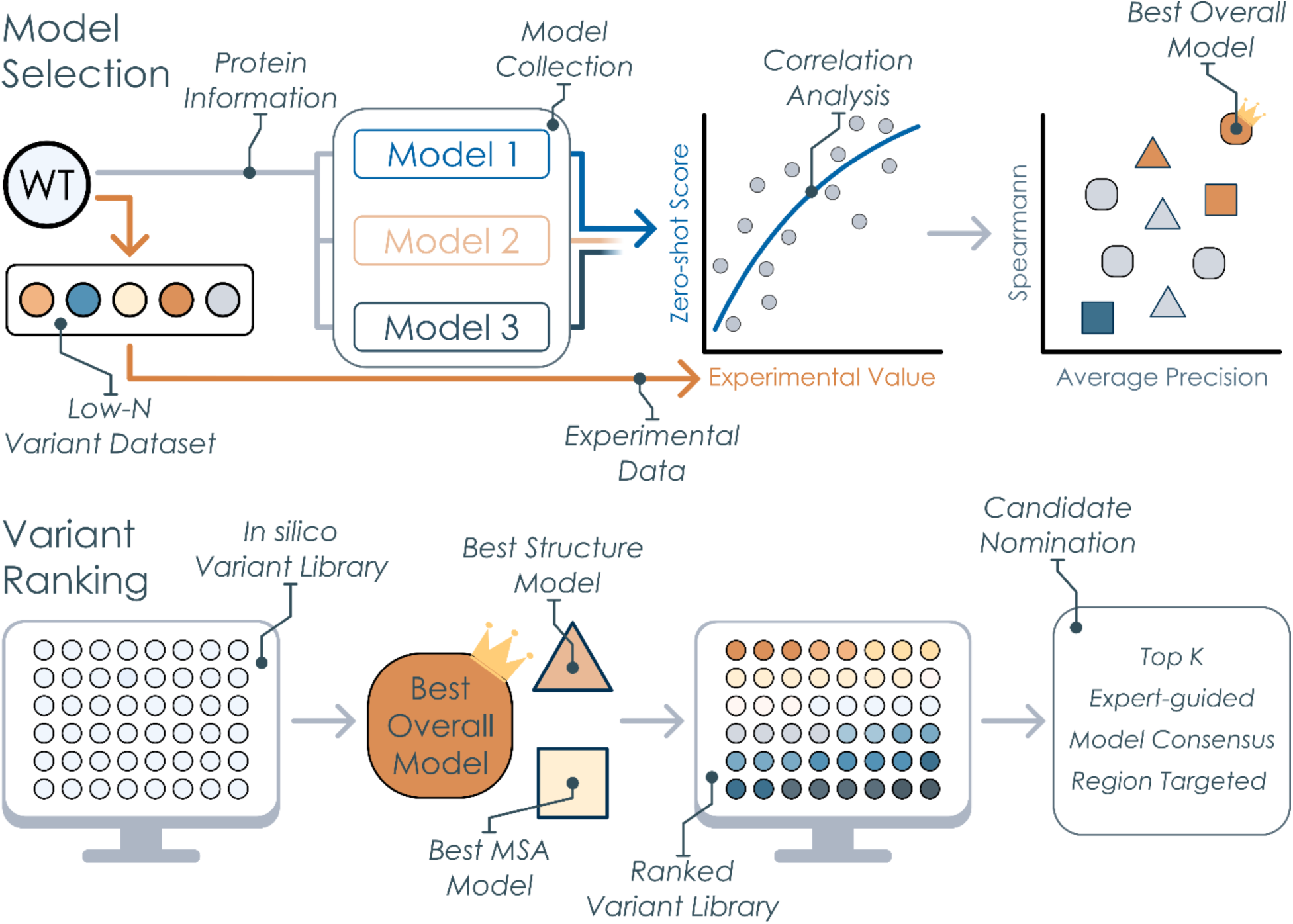
Overview of PRIZM workflow. The workflow consists of two phases: Model Selection and Variant Ranking. In the first phase, zero-shot predictors use the WT protein information to generate zero-shot scores for variants. These scores are compared to the experimental values in a low-N dataset of variants to identify the best-performing models. In the second phase, these models are applied to an in silico variant library to rank and nominate candidates for experimental characterization.

First, we validated the performance of PRIZM across ten literature benchmark datasets with a diverse set of protein properties. For all datasets except one, we found the workflow to consistently differentiate between models with high or low correlation to the assayed properties using as few as 20 variants, although 50 variants were generally preferred for identifying the best-performing models.

We further evaluated the workflow in two additional protein systems: sucrose synthase (SuSy) thermostability and UDP-dependent glycosyltransferase (GT1) activity. In the first case study, we utilized an in-house pre-existing dataset of 68 variants from a semi-rational engineering campaign as the low-N dataset for model selection (Mejia-Otalvaro *et al*. 2026). Based on a combination of the three best models identified by PRIZM, we identified a variant with an apparent melting temperature increase (ΔT_m, app_) of ∼3.0°C. In the GT1 activity study, the low-N dataset consisted of only eight variants from a previous rational campaign (Lax *et al*. 2026). By using zero-shot predictions to guide variant prioritization, we identified a variant with ∼20% increase in relative activity, with an overall hit rate of 60% for identifying variants with enhanced relative activity compared to WT.

Overall, our study highlights how PRIZM facilitates easy and data-efficient engineering of proteins while requiring little computational setup. We believe PRIZM can guide non-experts in utilizing zero-shot predictions for candidate selection, as well as provide experts with an informed choice of foundation models for downstream modelling.

## Materials and Methods

### PRIZM Workflow

PRIZM consists of two phases: the Model Selection phase, where the best zero-shot model is identified, and the Variant Ranking phase, where the best model is used to predict and rank an *in silico* library (Figure 1). As the first step of the Model Selection phase, we collect the protein information required as inputs for the pre-trained zero-shot models: the sequence, structure, and MSA. The sequences of all variants in the low-N dataset are obtained from the input file. If no crystal structure without gaps is available, a structure is predicted by tools such as Alphafold3 (Abramson *et al*. 2024) using the WT sequence. The WT sequence is also utilized to construct an MSA using the EVcouplings pipeline (Hopf *et al*. 2019), though a custom MSA can be employed if it has the correct format (A2M). Importantly, the resulting structure and MSA are used as zero-shot model inputs for all variants, not just WT.

No experimental data is used for finetuning or influencing the models. The collection of models is based on the work done by Notin et al. in ProteinGym (Notin *et al*. 2023). The current version of PRIZM consists of 25 models in total – nine models using sequence as input, six MSA-based models, eight models using structural inputs, and two models using all three types of information (Table S1). Importantly, PRIZM is not limited to only the models utilized in this study but can be extended to incorporate any newly released zero-shot model. For a thorough description of the individual models, we refer the reader to the original papers of the models and the description available at ProteinGym.

PRIZM determines the relationship between the output set of zero-shot scores from each model and the actual experimental values. Absolute Spearman correlation is used to examine the ranking performance of each model across the full range of experimental values. Absolute rather than signed correlation is used, as an anti-correlated model is equally informative for variant ranking. Then, a user-defined threshold – such as the WT value – is used to binarize the experimental values and calculate the average precision, which specifically evaluates the model’s ability to classify high-performing variants above the threshold. Both performance metrics are normalized and added together, resulting in a single performance metric for each zero-shot model. The metric describes how well each model can explain the experimental property and is used for identifying the best and worst models for the given low-N dataset. For users who wish to restrict model selection to correctly signed correlated models or to substitute alternative performance metrics entirely, these options are available within the PRIZM workflow. The sensitivity of model ranking to the specific scoring function design of PRIZM has not been formally evaluated. However, the availability of alternative metrics within the workflow allows users to assess this directly for their specific system. Importantly, PRIZM not only outputs the highest- and lowest-performing models overall, but also the best and worst models using either sequence, MSA, protein structure, or all three information types. This allows the selection of multiple high-performing models based on different protein information, a strategy shown here to result in enhanced variants.

In the second phase of PRIZM, the Variant Ranking phase, an *in silico* library is created by mutating the WT sequence. PRIZM allows specifying which regions to mutate, what residues to use, and how many substitutions to conduct, with the default setup being a full single-point mutagenesis of the entire WT protein. The resulting library is processed by the best models, and the zero-shot scores are used to rank all variants in the library, using the original correlation analysis of the low-N dataset. The final ranked *in silico* library can be utilized as a tool for identifying new variants using either a greedy *Top K* selection or by employing a targeted approach where ranking and expert knowledge are combined.

### Validation of PRIZM

PRIZM is a model-selection framework rather than a supervised predictor trained on experimental data. The zero-shot models are applied directly, without fine-tuning or parameter optimization, and the role of the experimental low-N dataset is therefore not to train a new model but to identify which pre-trained predictor best captures the target property. Conventional train/test splitting is thus not directly applicable. Instead, to assess whether PRIZM can identify high-performing predictors from limited labelled data, we used a subsampling procedure in which low-N datasets of varying sizes were repeatedly drawn from full DMS datasets. These low-N subsets were used only for model selection, after which the selected models were evaluated on the corresponding full DMS datasets. This design therefore assesses the robustness of PRIZM as a model-selection framework, rather than model generalization in the supervised learning sense.

To obtain DMS benchmark datasets for the PRIZM validation, we selected ten datasets from the ProteinGym dataset collection (Notin *et al*. 2023), covering a diverse range of protein properties and measurement modalities, including protein aggregation (Seuma, Lehner, and Bolognesi 2022), receptor activity (Jones *et al*. 2020), thermostability (Nutschel *et al*. 2020), inhibitor resistance (Brenan *et al*. 2016), fluorescence (Ellis *et al*. 2024), receptor binding (Starr *et al*. 2020), peptide binding (Araya *et al*. 2012), and enzyme activity (Romero, Tran, and Abate 2015, Chiasson *et al*. 2020, Hobbs *et al*. 2022) (Table S2). The seven non-enzyme datasets were chosen to collectively span distinct property types, while three datasets measuring enzyme activity were included to ensure coverage of catalytic function, a primary target in protein engineering. The only exclusion criterion applied was that datasets had to contain single-point variants, excluding datasets consisting exclusively of multi-point mutations. It should be noted that we leveraged the corresponding MSA and structure files described in ProteinGym for PRIZM inputs. From each full DMS dataset, we generated low-N subsets by subsampling between 10 and 200 variants, repeating this subsampling procedure 100 times (Figure S1). Both the full datasets and the corresponding low-N subsets were processed with PRIZM to identify high- and low-performing models. For each low-N dataset size, the models identified as best or worst by PRIZM were compared with their corresponding performance on the full dataset, and average performance across the 100 subsampling iterations was computed.

To compare the difference in performance of PRIZM between the best- and worst-performing models across the low-N dataset sizes, we calculated Cohen’s *d*, the standardized effect size:

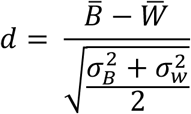

Where *B̅*, *σ*_*B*_, *W̅*, and *σ_W_* are the means and standard deviations of the performances of the best and worst models, respectively, across the 100 iterations (Cohen 1988). Following Cohen, we interpret a *d*-value of below 0.2 to be a small difference, between 0.2 and 0.5 to be a medium difference, and above 0.5 to be a large difference.

To assess the effect of mutation count on the PRIZM performance, we separated the protein aggregation (Seuma, Lehner, and Bolognesi 2022) and peptide binding (Araya *et al*. 2012) benchmark datasets into purely single or double mutant subsets. From these, we generated corresponding low-N datasets, processed them with PRIZM, and evaluated the best- and worst-performing models on the full datasets. This resulted in four comparisons: (i) performance on the full single-mutant dataset using models selected from single-mutant low-N data; (ii) performance on the full single-mutant dataset using models selected from double-mutant low-N data; (iii) performance on the full double-mutant dataset using models selected from single-mutant low-N data; and (iv) performance on the full double-mutant dataset using models selected from double-mutant low-N data. We again utilized Cohen’s *d* to quantify the difference between the best- and worst-performing models.

To compare PRIZM’s performance to the consensus approach of Hie et al. (Hie *et al*. 2024), we replicated their strategy on the benchmark datasets in this study. Using ESM-1b and the five ESM-1v versions, we identified variants that were predicted to be better than WT for multiple of these models. As described by Hie et al., we adjusted the consensus model count to limit how many variants were nominated, iteratively increasing the number of models until fewer than 40 variants were proposed. Furthermore, we implemented a similar consensus approach for the top-performing PRIZM models across the four protein information types: sequence, MSA, structure, and all. These two consensus approaches were compared to a simple greedy top 10 approach leveraging the best-performing PRIZM model. For each approach, we calculated the hit rate, with the performance of the two PRIZM-based strategies being based on 100 iterations using a low-N dataset of 50 variants.

### Generation of PRIZM inputs for Case Study proteins

The MSAs of *Gm*SuSy and TOGT1_1 were created using EVcouplings (Hopf *et al*. 2019) with a bitscore cutoff of 0.5 and 0.5, respectively. For *Gm*SuSy, both a monomeric and a tetrameric structure were generated using AlphaFold3 (Abramson *et al*. 2024), while we only generated a monomeric structure of TOGT1_1. Only the monomeric structures were utilized as input for PRIZM. To obtain the *Gm*SuSy low-N dataset, we subsampled all single mutants from the consensus mutagenesis campaign described in Mejia-Otalvaro et al. (Mejia-Otalvaro *et al*. 2026). The TOGT1_1 low-N dataset contained all single-mutant substitutions described by Lax et al. (Lax *et al*. 2026).

### Nomination of variants for experimental validation in Case Studies

To nominate new *Gm*SuSy variants, we ranked every single-mutant using Tranception No Retrieval, MIFST, and MSA Transformer. These ranks were added together, resulting in a composite ranking of each variant. This was used to select the top 5 best variants, from which 3 had already been characterized in the original low-N dataset. We therefore selected the last two, L731E and F468I, for experimental characterization.

To nominate new TOGT1_1 variants, we constructed a full single-mutant landscape using the VenusREM scores, identifying six positions with high predicted mutability: 53, 72, 208, 239, 389, and 401. At each position, we examined the TOGT1_1 structure to identify any steric clashes caused by the mutations. For all positions except 53, the highest-ranked mutation was chosen as no steric clashes were predicted by expert analysis. However, the best variant at 53, N53Y, showed potential for structural conflicts, and we therefore chose the second variant. Lastly, we selected the third-ranked variant at position 401, G401F, as i) the site appeared able to accommodate a bulkier residue, and ii) the Gly to Phe substitution introduced a substantial local change, making it an interesting mutation to test. Our final nomination of new TOGT1_1 variants for experimental characterization was N53A, N72S, D208L, W239I, E389A, G401I, and G401F.

### Experimental validation

The experimental assays and materials used in the two case studies follow the original publications by Mejia-Otalvaro et al. (Mejia-Otalvaro *et al*. 2026) and Lax et al. (Lax *et al*. 2026). A summary of the experimental procedures is provided in Supplementary Methods.

### Computational Setup

All computations were performed on a Microsoft Azure Standard NC6s v3 virtual machine (6 vCPUs, 112 GiB RAM, NVIDIA Tesla V100 16 GB GPU, Ubuntu 24.04 LTS). Most models utilize GPU acceleration, which is the primary computational bottleneck, while a subset of models run on CPU. Runtimes vary substantially across models, with the fastest models scoring a full single-mutant library within minutes and slower models requiring several hours. The most time-intensive step is the “evotuning” of UniRep, which can require approximately one day depending on MSA depth. Overall, the computational requirements are modest by modern standards and accessible on standard cloud computing infrastructure.

## Results

### Validation of PRIZM

To examine the performance of PRIZM, we validated the workflow on benchmark deep mutational scan (DMS) datasets. Except for the inhibitor resistance of Mitogen-activated protein kinase 1 (MAPK1) (Brenan *et al*. 2016), we observed a Cohen’s *d* > 0.5 for the separation between best and worst models using datasets containing as few as 20 variants (30 for enzyme activity of epoxide reductase (Chiasson *et al*. 2020)). As *d* > 0.5 denotes a large difference, this suggests that ∼20 variants are sufficient for PRIZM to reliably distinguish between high- and low-performing models (Figure 2A, Figure S2). Datasets below this threshold may yield unreliable estimates of Spearman correlation and average precision, as both metrics exhibit substantially higher variance when estimated from fewer than ∼20 variants (Figure S3-4). Moreover, using ∼50 variants generally led to models with performance similar to the best overall model, and further increases in the labelled dataset size used for model selection yielded only minor improvements. Model ranking was found to be robust across a range of binarization thresholds from the 20th to the 80th percentile of the experimental distribution, for both the full dataset and subsamples of 50 variants (Figure S5).

**Figure 2.**
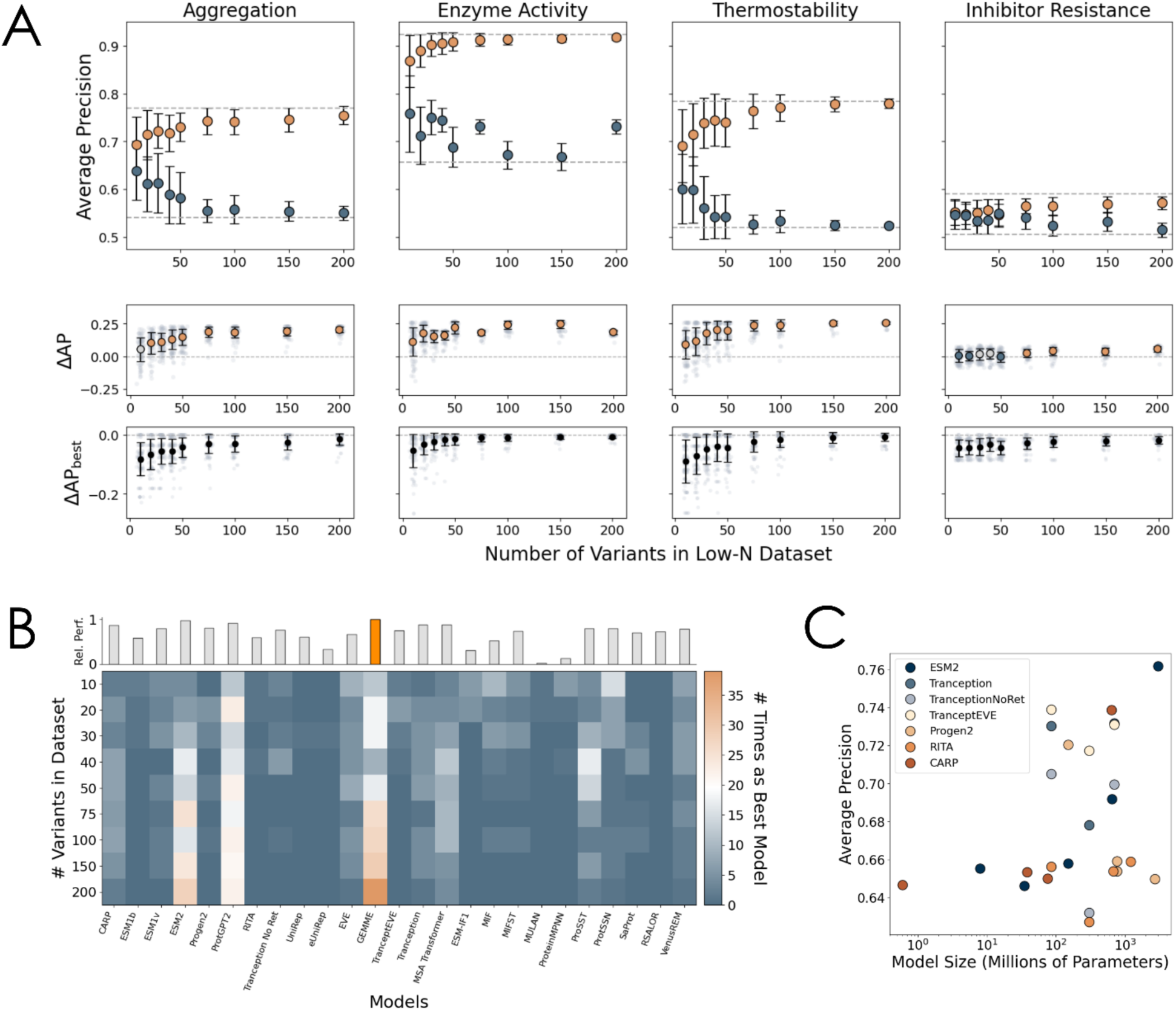
Validation of PRIZM workflow. **A** Performance of zero-shot models selected by PRIZM for predicting full DMS datasets as a function of the number of variants in the low-N dataset (aggregation data from Seuma et al. (Seuma, Lehner, and Bolognesi 2022), enzyme activity from Hobbs et al. (Hobbs et al. 2022), thermostability from Nutschel et al. (Nutschel et al. 2020), inhibitor resistance from Brenan et al. (Brenan et al. 2016)). The top row shows average precision (AP) for the best (orange) and worst (blue) models as identified by PRIZM, with dotted lines indicating the overall best and worst models. The middle row shows ΔAP between the best and worst models, colored by Cohen’s d (blue: d < 0.2, grey: d < 0.5, orange: d > 0.5). The bottom row shows ΔAP between each selected model and the best overall model (ΔAP_best_). Error bars represent the standard deviation from 100 PRIZM iterations. **B** Frequency of each model being selected as the best by PRIZM for different low-N dataset sizes for the aggregation DMS dataset from Seuma et al. (Seuma, Lehner, and Bolognesi 2022) across 100 iterations. The top inset shows each model’s relative performance by comparing its average precision with that of the overall best model (orange) on the full dataset. **C** Average model precision as a function of model size for the aggregation DMS dataset from Seuma et al. (Seuma, Lehner, and Bolognesi 2022).

We further examined how often the models with the highest overall performance of the full DMS datasets were chosen by PRIZM as a function of the low-N dataset size (Figure 2B, Figure S6-14). Here, we found that PRIZM generally converged towards high-performing models and successfully identified the best models for most of the benchmark datasets. The identity of the top-performing model also varied across datasets, underscoring that no single model consistently dominated performance. Furthermore, consistent with recent analyses suggesting diminishing returns from model scaling (Pascal Notin 2025), we observed that increasing the number of model parameters did not necessarily lead to improved performance (Figure 2C, Figure S15). Together, our analysis highlights not only the general accuracy of PRIZM for selecting good models but also the importance of the Model Selection Phase as a whole.

Next, we assessed the impact of mutation count per variant by evaluating PRIZM on datasets restricted to single or double mutants (Figure S16-17). We found a general decrease in performance of the zero-shot models when predicting the effects of double mutants, which could potentially be caused by the models’ poor understanding of epistatic effects. PRIZM’s ability to distinguish between high- and low-performance models also appears to weaken in the case of learning from double mutants for datasets with fewer variants.

Lastly, to compare the performance of PRIZM to a previous zero-shot prediction workflow, we compared PRIZM with the approach described by Hie et al. (Hie *et al*. 2024). First, we applied their approach to the benchmark datasets described in this study. Secondly, we assessed the hit rate of PRIZM of the top 10 variants ranked by the top-performing model. As a last step, we adapted the consensus approach of Hie et al., using the best models of each protein information type. PRIZM outperformed the approach of Hie et al. (Hie *et al*. 2024) across six of the ten benchmark datasets (Figure S18). Furthermore, a key limitation of the Hie et al. approach is that it can only propose mutations that are predicted to outperform WT by multiple zero-shot models simultaneously. If no such variants exist for a given protein, the method cannot nominate any candidates. On the contrary, PRIZM always provides a ranked *in silico* library of any chosen model. This gives researchers full flexibility in how they prioritize mutations, whether by employing a greedy *Top K* approach, combining multiple model rankings, or incorporating expert domain knowledge.

In summary, our validation highlights the versatility and generalizability of PRIZM for model selection and variant prioritization. The approach extends beyond enzyme activity to protein properties such as thermostability, receptor activity, and fluorescence, reliably distinguishing high- and low-performing models with as few as 20 variants, and as few as 50 variants are usually enough to reach the upper limit on the performance, based on the retrospective analysis of published datasets. The method can handle double mutants as well, although a lower average precision is achieved when predicting the effects of double-point mutations. Overall, PRIZM provides a scalable, property-agnostic strategy for navigating the space of zero-shot models while benefiting from available small experimental datasets.

To demonstrate how PRIZM can help guide protein engineering campaigns in real applications, we next engineered the thermostability of sucrose synthase and catalytic activity of a glycosyltransferase.

### Case Study 1 – Improving the thermostability of sucrose synthase using an existing dataset

As mentioned previously, state-of-the-art MLDE approaches often emphasize the creation of new, unbiased datasets (Jiang *et al*. 2025). Here, we evaluated PRIZM’s ability to utilize existing datasets, thereby minimizing the requirement for new experimental characterization.

As a case study, we investigated the thermostability of sucrose synthase from *Glycine max* (*Gm*SuSy). SuSy is able to catalyze the reversible transfer of glucose from sucrose to nucleotide diphosphates (typically UDP), forming UDP-glucose and fructose. Owing to this activity, SuSy enzymes are frequently employed in UDP-glucose regeneration systems for GT1-coupled reactions (Mejia-Otalvaro *et al*. 2025).

We have previously shown how the engineering of *Gm*SuSy thermostability can be guided using tools such as ancestral sequence reconstruction and consensus mutagenesis (Mejia-Otalvaro *et al*. 2026). Based on the latter, we obtained T_m, app_ dataset consisting of single mutants distributed across the full protein sequence and their melting temperature (T_m, app_) (Figure 3A). This dataset was used as input for PRIZM to identify the most suitable zero-shot model for predicting T_m, app_ (Figure 3B, Figure S19-20). Among the top-performing models, two demonstrated comparable performance: Tranception No Retrieval and MIFST, which rely primarily on sequence and structural information, respectively. We selected both models along with MSA Transformer, the second-best-performing MSA-based model. We did not choose the best MSA-based model, Tranception, to avoid potential redundancy with two Tranception-based models.

**Figure 3.**
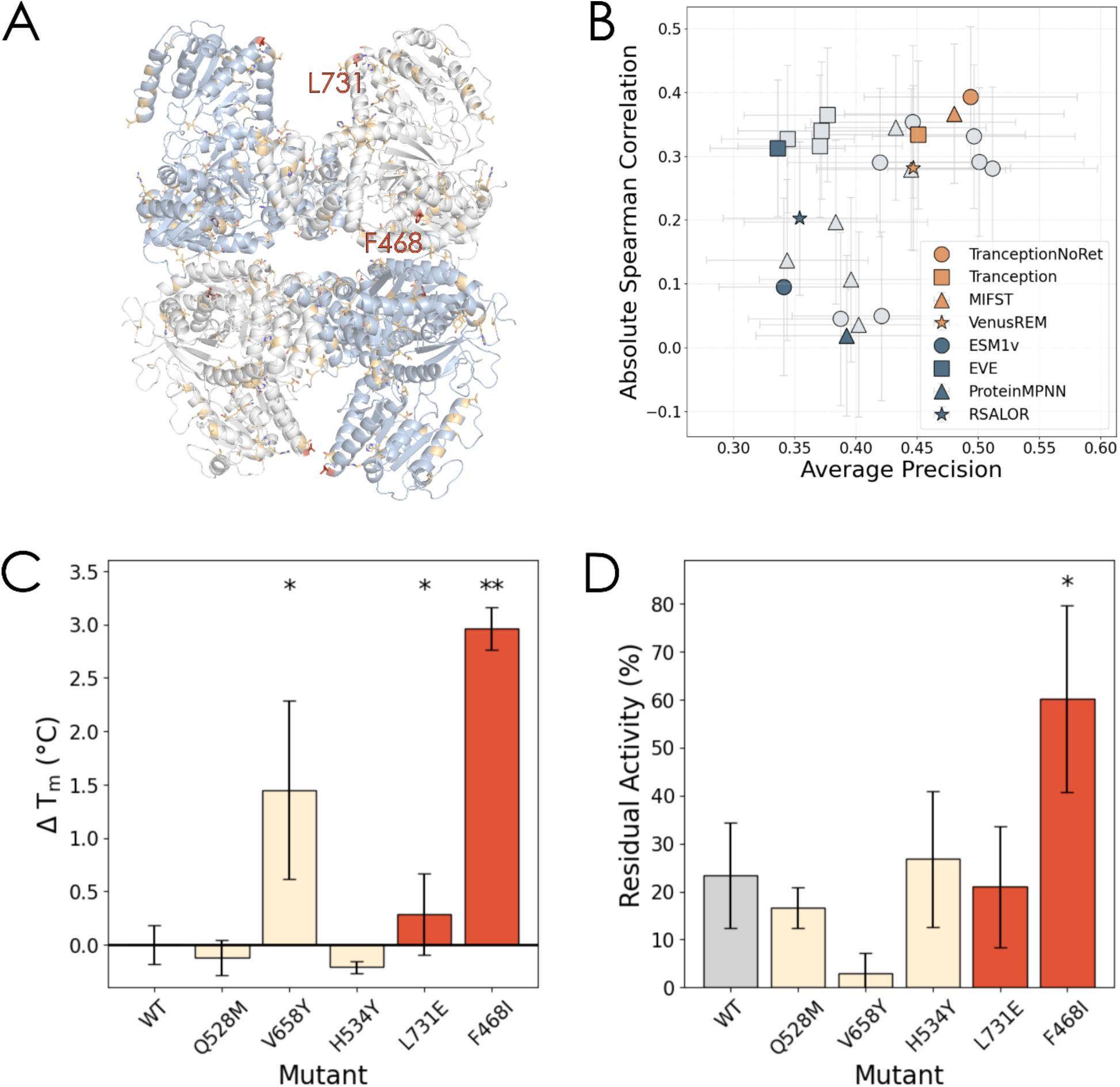
PRIZM engineering of GmSuSy. **A** A predicted structure of the GmSuSy tetramer using AlphaFold3. Positions mutated in the original consensus mutagenesis campaign by Mejia-Otalvaro et al. (Mejia-Otalvaro et al. 2026) are shown in orange, while the positions mutated in this study are shown in red. **B** Spearman correlation and average precision of the PRIZM zero-shot models for predicting GmSuSy T_m, app_. Circle = sequence-based model, square = MSA-based model, triangle = structure-based model, star = all three. Blue and orange marker colours denote the lowest and highest performing models for each model category, respectively, as identified by PRIZM. Error bars indicate the standard deviation evaluated from bootstrapping the initial dataset (N = 1000). **C, D** The experimental ΔT_m, app_ (**C**) and residual activity after 5 min incubation at 60°C (**D**) of WT GmSuSy and the top seven variants predicted to be better than WT by Tranception No Retrieval, MSA Transformer, and MIFST, sorted by the combined rank. Grey = WT, beige = variants from previous campaign, red = new variants identified by PRIZM. Asterisks denote relative activities above WT (one-sided Welch test with α = 0.05 or α = 0.01, indicated by one or two asterisks, respectively), while the error bars indicate error propagated replicate standard deviation (N = 3).

We then processed an *in silico* library of all possible single-point variants using the three selected models and identified the variants predicted to outperform WT across all models (Figure S21). Notably, the resulting list included several variants that were already present in the original rationally designed dataset, showing a convergence between expert-driven rational mutagenesis and zero-shot predictions, even though the algorithm is naïve to said expert-driven mutagenesis. When ranking variants by their combined performance across the three models, these previously characterized mutants occupied three of the top five positions, with the standout variant V658Y having a ΔT_m, app_ of ∼1.7°C (Figure 3C, Table S3). We further characterized the remaining two variants, absent from the original dataset, L731E and F468I, which both had higher ΔT_m, app_ than WT (∼0.3°C and ∼3.0°C, respectively). Overall, while moderate in the absolute numbers, our PRIZM analysis achieved a 60% hit rate of variants with higher T_m, app_ than WT. Furthermore, the best variant, F468I, outperformed all variants in the original dataset (Figure 3C, Table S3). This increase in melting temperature was accompanied by kinetic stability towards elevated temperature, as supported by residual activity assays at 60°C, where F468I retained over 60% activity, compared to ∼23% for WT (Figure 3D).

### Case Study 2 – Engineering a glycosyltransferase using fewer than ten variants as initial dataset

To assess the robustness of PRIZM’s Model Selection Phase, we evaluated its performance in an extremely low-N setting. In our prior work, we conducted an engineering campaign of the GT1 from *Nicotiana attenuata* (TOGT1_1) (Lax *et al*. 2026). GT1s catalyze the transfer of sugar moieties to a diverse set of acceptor molecules, offering a biocatalytic solution to the synthesis of industrially relevant glycosylated compounds (Gharabli, Della Gala, and Welner 2023, Sirirungruang et al. 2023). We focused on niclosamide, an essential small molecule therapeutic that suffers from low water solubility, which limits its therapeutic use beyond tapeworm treatment. We previously identified TOGT1_1 as a niclosamide-active GT1 (Harding-Larsen, Madsen *et al*. 2024, Lax *et al*. 2026) and performed a small rational design campaign of eight single active site mutants to improve the catalytic efficiency, with the best mutant having 109% relative activity compared to WT.

We used this dataset of eight single mutants as input to PRIZM (Figure 4A), through which PRIZM determined VenusREM to be the top-performing model (Figure 4B, Figure S22-23). This model integrates all three protein information types: sequence, structure, and MSA.

**Figure 4.**
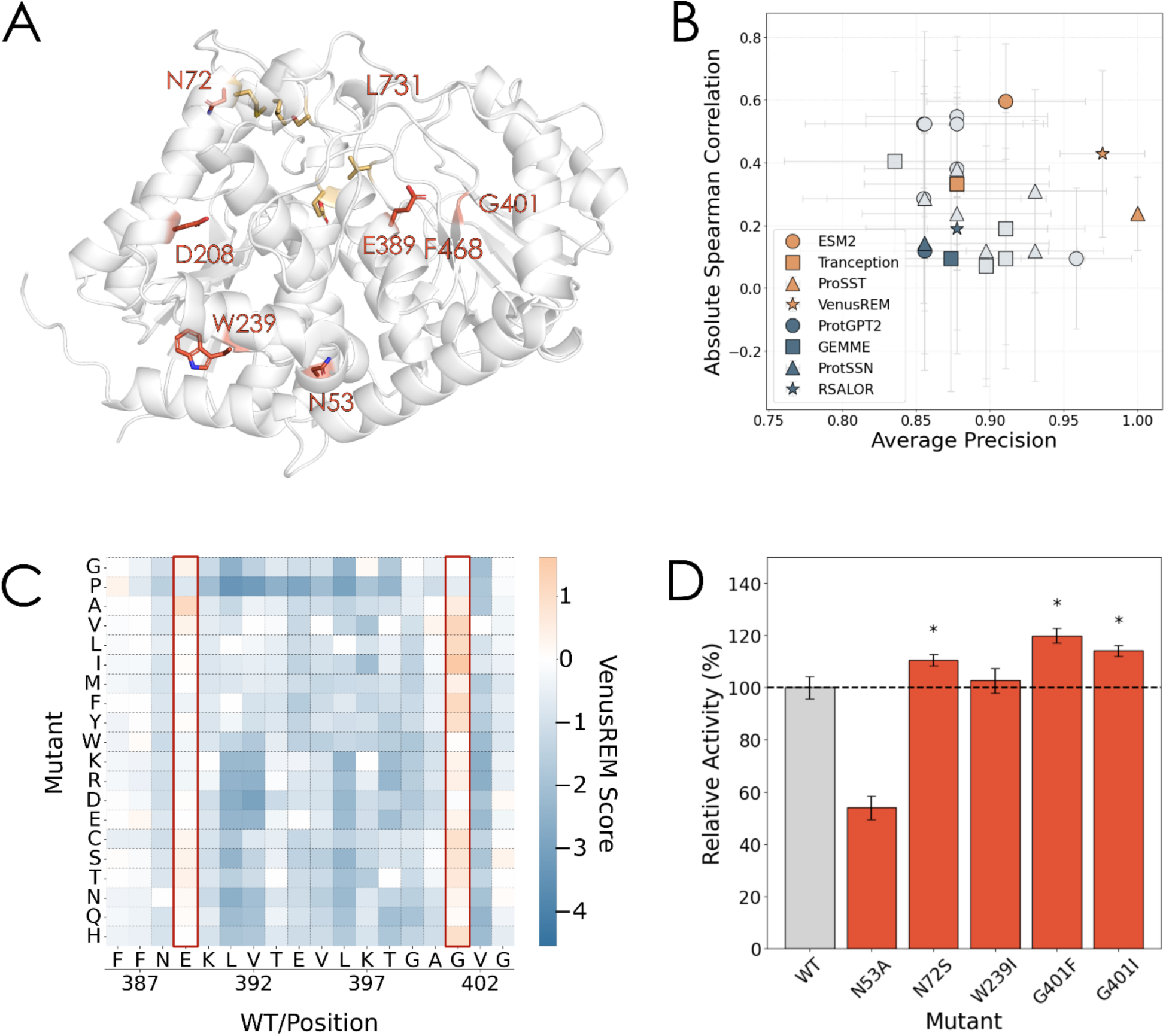
PRIZM engineering of TOGT1_1. **A** A predicted structure of TOGT1_1 using AlphaFold3. Positions mutated in the original rational design dataset from Lax et al. (Lax et al. 202C) are shown in beige, while the positions mutated in this study are shown in red. **B** Spearman correlation and average precision of the PRIZM zero-shot models for predicting TOGT1_1 relative activity. Circle = sequence-based model, square = MSA-based model, triangle = structure-based model, star = all three. Blue and orange denote the worst and best models for each model category, respectively, as identified by PRIZM. Error bars indicate the standard deviation using a bootstrapping approach (N = 1000). **C** Representative section of TOGT1_1 in silico landscape of VenusREM scores. Orange denotes variants predicted to be better than WT, while blue denotes variants predicted to be worse. Red squares mark E28S and G401; selected for experimental characterization due to high proposed mutability. **D** Relative activities of TOGT1_1 variants selected using the TOGT1_1 VenusREM landscape. Asterisks denote relative activities above WT (one-sided Welch test with α = 0.05), while the error bars indicate replicate standard deviation (N = 3).

Similar to the previous PRIZM engineering campaigns, we used VenusREM to process an *in silico* library of all possible single mutants. Rather than directly selecting top-ranked substitutions, we analyzed the VenusREM mutational landscape to guide variant selection. Here, we identified positions with high apparent mutability, defined as those where VenusREM predicted the majority of substitutions to result in enhanced performance (Figure 4C, Figure S24). VenusREM predicted the positions of 53, 72, 208, 239, 389, and 401 to have high predicted mutability, and seven single mutants were selected for further characterization based on a combination of zero-shot ranking, chemical knowledge, and protein structure examination (Figure 4D, Table S4). For N72, D208, W239, and E389, we chose the highest-ranked variant, while we chose an Ala substitution instead of Trp for N53 due to apparent backbone restrictions. For G401, we selected both the highest-ranked variant and the third-highest-ranked variant, due to the latter representing a drastic biochemical change from Gly to Phe. We were unable to express two of these variants, but of the expressed mutants, we identified N53A, G401F, and G401I, with relative activities of 110.5%, 119.9%, and 114.1% compared to the WT, respectively. This corresponded to a hit rate of ∼60% for variants better than WT. While the observed improvements are small in magnitude, the best-performing variants, G401F and G401I, perform comparably to the best variants from the original engineering campaign. Furthermore, G401F and G401I are situated in a coil-rich region distant from the active site, which would likely be overlooked in a rational design campaign. Interestingly, the best variant, G401F, was ranked as the 3^rd^ best G401 substitution (11^th^ best overall) by VenusREM. This underscores the strength of combining zero-shot predictors and expert evaluation for exploring the protein sequence space.

## Discussion

This study describes PRIZM, a two-phase approach for leveraging pre-trained zero-shot models to predict the effects of protein variants, offering a robust and accessible protein engineering workflow for non-experts. By identifying the zero-shot models that best describe a target protein property, PRIZM combines the simplicity of zero-shot modelling with the task-specific insight typically associated with supervised learning. In doing so, it addresses two major challenges: the difficulty of supervised ML in low-N settings and the lack of guidance in standard zero-shot modelling, all without requiring model training, dataset construction, or specialized ML knowledge.

Beyond this, PRIZM represents an orthogonal strategy to supervised learning as it focuses on the selection of a suitable pre-trained model rather than its downstream application. Often, this selection is either arbitrary or based on a global validation across multiple benchmark datasets. PRIZM enables researchers to generate zero-shot scores and protein embeddings that are meaningful, property-specific, and enriched with protein-relevant biological knowledge.

In practice, PRIZM is particularly useful in three scenarios: when a small mutational dataset from a previous engineering campaign is already available, when experimental budget constraints limit the feasibility of multiple iterative design rounds, and when researchers aim to leverage zero-shot predictors without extensive ML expertise. Here, our benchmarking suggests that PRIZM can generally distinguish high- and low-performing models using ∼20 variants, while results below this regime should be interpreted with caution. Nonetheless, the TOGT1_1 case study indicates that even in very low-N settings, PRIZM may still provide useful guidance when combined with expert curation of the ranked candidates. Detailed guidance on how to use PRIZM can be found in Supplementary Methods.

We validated PRIZM across ten benchmark DMS datasets, finding that the models ranked as best by PRIZM consistently outperformed the lowest-ranked ones using as few as 20 variants; the best models chosen using only 50 variants ranked among the top performers from the full DMS datasets. We further conducted two case studies targeting the thermostability of *Gm*SuSy and the activity of TOGT1_1, illustrating the application of PRIZM to distinct engineering tasks across different target proteins and properties in contexts constrained by low-throughput experimental assays. Our validation and case studies also highlight the framework’s ability to leverage data from previous engineering campaigns, thereby opening new opportunities for researchers to repurpose existing experimental datasets for model-driven protein design.

Beyond serving as a tool for zero-shot-based protein engineering, PRIZM can also act as an initial step within a larger design pipeline, potentially mitigating some of the limitations of PRIZM. In particular, the quality of variant predictions is inherently constrained by the accuracy of the best-performing zero-shot model. As highlighted when processing the inhibitor resistance dataset of MAPK1 (Brenan *et al*. 2016), PRIZM may struggle with properties for which no pre-trained model adequately captures the relevant mutational effects. In the case of MAPK1, the inhibitor resistance was found to be uncorrelated with protein function. Zero-shot models learn from evolutionary sequence variation and therefore capture functional constraint well, but carry no signal for resistance to a synthetic inhibitor that was absent from evolutionary history. As such, zero-shot predictions are expected to struggle with properties governed by protein-drug interactions, regulatory evasion, or other context-specific mechanisms that leave no imprint on sequence evolution. Additionally, the ranking of double mutant libraries revealed potential challenges with modelling epistatic interactions. These limitations could be addressed by integrating PRIZM into a supervised framework that leverages the best zero-shot model, such as substituting the ESM2 embeddings of the EVOLVEpro workflow with those from the top-performing embedding model (Jiang *et al*. 2025), or augmenting the zero-shot score with a one-hot encoding as described by Hsu et al. (Hsu, Nisonoff *et al*. 2022).

Another key challenge when using PRIZM lies in selecting variant candidates for further characterization. Relying solely on a greedy *Top K* selection strategy limits the sequence exploration, while incorporating expert knowledge risks diminishing the contribution of the latent biological patterns captured by large foundation models. A potential solution could involve employing Bayesian optimization, as proposed by Hie et al. (Hie and Yang 2022), where the model uncertainty guides the selection of new variants.

Collectively, these considerations underscore PRIZM’s potential to serve as a foundation for bridging zero-shot modelling and data-driven protein design. We envision PRIZM as a resource to inform and accelerate future protein engineering efforts for experts and non-experts alike, whether employed as an independent predictive tool or integrated within larger design pipelines.

## Supporting information

GmSuSy variant zero-shot scores and experimental results

TOGT1_1 variant zero-shot scores and experimental results

## Data Availability Statement

All code and datasets used to produce the results and figures in this manuscript are publicly available via Zenodo (PRIZM v1.1.1; DOI: 10.5281/zenodo.19483130) and the PRIZM GitHub repository (release v1.1.1; https://github.com/daha-la/PRIZM). The GitHub repository also includes documentation and installation/running instructions. The archived Zenodo release contains the complete codebase, analysis scripts used to reproduce the figures, and the datasets underlying this study.

## Funding

This work was supported by the Novo Nordisk Foundation via grants no. NNF20CC0035580 (The Novo Nordisk Foundation Center for Biosustainability), NNF24SA0100980 (The Novo Nordisk Foundation Biotechnology Research Institute for the Green Transition), and NNF23OC0085634 (B.M.L.). Additional funding was provided by the Czech Ministry of Education, Youth and Sports (ESFRI RECETOX RI LM2023069, ESFRI ELIXIR LM2023055, and e-INFRA CZ 90254) as well as the European Union’s Horizon 2020 Research and Innovation Programme through grant agreement No. 857560 (CETOCOEN Excellence) and the European Union’s Horizon Europe Framework Programme under the grant agreement No. 101136607 (CLARA). This publication reflects only the author’s view, and the European Commission is not responsible for any use that may be made of the information it contains. Felipe Mejia-Otalvaro received support from the Copenhagen Bioscience Ph.D. Programme (Grant No. NNF22SA0078231).

## Acknowledgements

Parts of the thermostability assays were conducted during the MSc course, 29902 – Advanced Synthetic Biology for Creating Sustainable Cell Factories at the Technical University of Denmark, and we thank Anne Sofie Gadfelt, Emil Victor Solvang Lundquist, and Xizhi Hong for their contributions. We also thank Karen Pailozian for feedback regarding the accessibility of the PRIZM GitHub repository. AI-assisted tools were used to support manuscript preparation and code development. Specifically, AI-tools were used for grammar and phrasing suggestions and to assist with drafting and debugging the code. All content and code were reviewed, edited, and validated by the authors, who take full responsibility for the final manuscript and software.

## Conflicts of Interest

The authors have no conflicts of interest to declare.

## Notes

### Competing Interest Statement

The authors have declared no competing interest.

### Summary of Updates

Manuscript reorganized to Introduction/Materials and Methods/Results/Discussion; Validation of PRIZM section updated to clarify the model-selection evaluation framework; Discussion revised and practical guidance moved to the Supplementary Information; figures and supplementary files updated; data availability updated to PRIZM v1.1.1 / new Zenodo archive.

