## Supplementary Figures and Tables for "PRIZM: Combining Low-N Data and Zero-shot Models to Design Enhanced Protein Variants"

### Supplementary Methods

#### Case study 1

All materials and methods for Case Study 1, including protein expression, purification, and enzyme assays, were performed as described in Mejia-Otalvaro et al. (Mejia-Otalvaro *et al.* 2026), with identical buffer compositions and instruments. A brief summary is provided below.

**Cloning and transformation**

*Gm*SuSy-WT was obtained in a pET28a(+) expression vector containing an N-terminal His-tag and TEV protease cleavage site. Amino acid substitutions were introduced by PCR-based cloning, and all constructs were sequence-verified by Sanger sequencing prior to expression. Confirmed plasmids were transformed into chemically competent *E. coli* BL21 Star (DE3) cells.

**Protein expression and purification**

Expression strains were cultivated in kanamycin-supplemented rich medium at 37°C until an optical density at 600 nm ($OD_{600}$) between 0.5 and 0.8, followed by induction with isopropyl-β-D-thiogalactopyranoside (IPTG) and overnight protein expression at reduced temperature. Cells were harvested by centrifugation, resuspended in lysis buffer, and disrupted by sonication under cooling conditions. Clarified lysates were purified by $Ni^{2+}$-affinity chromatography using a HisTrap FF column on an ÄKTA pure system (Cytiva Life Sciences). Target protein fractions were identified by SDS-PAGE, pooled, buffer-exchanged, concentrated using 50 kDa centrifugal filters, aliquoted, and stored at −70°C. Protein concentrations were determined by UV absorbance at 280 nm.

**Thermostability assays**

Protein thermodynamic stability was evaluated by determining apparent melting temperatures ($T_{m,app}$) using differential scanning fluorimetry (DSF) through Protein Thermal Shift Dye kit. Fluorescence-based unfolding profiles were recorded during controlled heating using a QuantStudio 5 real-time PCR system (Applied Biosystems). Measurements were performed with four technical replicates.

**Residual activity assays**

Residual enzymatic activity as a proxy for kinetic stability was assessed following short-term thermal treatment of protein samples prior to reaction initiation. Enzymatic activity was quantified using a sucrose/UDP-based assay, and fructose formation was measured with a commercial enzymatic detection kit (Megazyme). Residual activity was calculated relative to the activity of the thermal untreated wild-type enzyme and used as a measure of kinetic stability. All measurements were performed in triplicate.

#### Case study 2

All materials and methods for Case Study 2, including protein expression, purification, and enzyme assays, were performed as described in Lax et al. (Lax *et al.* 2026), with identical buffer compositions and instruments. A brief summary is provided below.

**Cloning and transformation**

The WT TOGT1_1 gene was previously ordered with in a pET28a(+) expression vector containing N- and C-terminal His-tags and a TEV protease cleavage site from Biomatik(Harding-Larsen *et al.* 2024). All amino acid substitutions introduced by PCR-based site-directed mutagenesis, and all constructs were sequence-verified by Sanger sequencing prior to expression. Confirmed plasmids were transformed into chemically competent *E. coli* BL21 Star (DE3) cells.

**Protein expression and purification**

Expression strains were cultivated in kanamycin-supplemented rich medium at 37°C until reaching an $OD_{600}$ of 0.8-1, followed by induction with 250 µM IPTG and overnight protein expression at reduced temperature. Cells were harvested by centrifugation, resuspended in lysis buffer, and disrupted by sonication under cooling conditions. Clarified lysates were purified by $Ni^{2+}$-affinity chromatography using a HisTrap FF column on an ÄKTA pure system (Cytiva Life Sciences). Target protein fractions were identified by SDS-PAGE, pooled, buffer-exchanged, concentrated using 50 kDa centrifugal filters, aliquoted, and stored at −70°C. Protein concentrations were determined by UV absorbance at 280 nm.

**Enzyme activity assays**

The TOGT1_1 glycosylation reactions were measured using purified protein (50 µg/mL), Nic/Nic-Glc (200 µM), and UDP-Glc (5 mM) in 100 µL reaction buffer and 20% (v/v) DMSO. All reactions were incubated at 30°C, quenched with equivolume methanol, filtered, and analyzed by reversed-phase HPLC (Ultimate 3000 Series HPLC, Thermo Fisher Scientific). HPLC analysis was performed using a ZORBAX RR Eclipse Plus C18 column (100 × 4.6 mm, 3.5 µm; Agilent) at 30°C. Mobile phases were water + 0.1% formic acid (A) and acetonitrile (B), with a 1 mL/min gradient elution following the method described in Lax et al.(Lax *et al.* 2026). Chromatograms were recorded at 330 nm using Chromeleon 7.2.9, and relative activities were determined from integrated peak areas.

#### Guidance for using PRIZM

Here, we provide practical guidance on when and how to implement PRIZM most effectively.

Regarding when to leverage the workflow for variant nomination, three settings generally favor the use of PRIZM. The first is the presence of a mutational dataset from prior engineering campaigns. Unlike several state-of-the-art methods (Zhou *et al.* 2024, Jiang *et al.* 2025, Teufel *et al.* 2025), PRIZM is designed to work with any existing mutational dataset, and we therefore encourage researchers to leverage PRIZM together with datasets from previous campaigns. The second setting favoring PRIZM is when budget restrictions allow for only a few rounds of experiments, as this will reduce the effectiveness of few-shot modelling approaches such as EVOLVEpro (Jiang *et al.* 2025). The PRIZM rankings do not depend on multiple rounds of experiments and can thus be utilized as a single-round nominator. Lastly, researchers with limited ML expertise might also benefit from using PRIZM instead of more elaborate supervised strategies that require model fine-tuning, hyperparameter optimization, and careful train-test splitting. Conversely, when no prior experimental data exists and budget allows for multiple iterative rounds, supervised or few-shot approaches are likely to achieve higher predictive accuracy and may be preferred. Here, we suggest using PRIZM to identify the best initial model for subsequent fine-tuning.

When running PRIZM, it is important to evaluate the performance metrics of the best models, as the random baseline of these metrics are influenced by both dataset size and bias (Figure S21-22). If the best model is worse than a random baseline, its predictions may not reliably capture mutational effects. As PRIZM can never be better than the best model, we do not advise the use of PRIZM in such a scenario.

When choosing a threshold value for defining low and high-performing variants, we advise using the WT value, as this will naturally divide the data into variants better and worse than the WT. However, as the PRIZM model rankings were found to be robust across a range of binarization thresholds, the specific threshold should not substantially alter the general rankings (Figure S5).

Our retrospective DMS benchmarking demonstrated that PRIZM can robustly distinguish between high- and low-performing models using as few as 20 variants. For any dataset sizes below this, we advise caution when utilizing the workflow. Although PRIZM successfully identified improved TOGT1_1 variants using only 8 initial mutants, we incorporated expert evaluation in addition to the model rankings. An expert curation of the PRIZM results can thus be a potential avenue for utilizing the workflow in very low-N settings.

For low-N datasets of 20 or more variants, the best models identified by PRIZM generally provide reliable guidance for selecting new variants. If one model clearly outperforms all other models, we recommend using only this model for ranking the *in silico* library. However, if several models exhibit similar performance, our results suggest utilizing multiple models for variant selection. This can either be implemented using a simple additive combination of the variant ranks, as demonstrated in our *Gm*SuSy case study, or by more elaborate ensemble strategies such as that proposed by Hie et al. (Hie *et al.* 2024). We suggest mainly combining models based on different protein information types, to reduce the potential for redundant information.

Finally, when selecting the variants for further experimental characterization, we recommend balancing exploitation (testing the highest-ranked candidates) with exploration (sampling diverse regions of sequence space). Although some substitutions may appear nonsensical from the perspective of domain expertise, discarding such variants outright contradicts the motivation for using zero-shot models, which is to identify beneficial mutations that lie outside conventional domain knowledge. Conversely, some model rankings may be dominated by mutations at a single position and prioritizing only these can result in a narrow search of the mutational landscape. To avoid this, we suggest selecting the top-ranked variants while ensuring coverage across multiple positions. If there are a large number of positions with high apparent mutability, the model rankings can also be used for the design of smart libraries.

#### Supplementary Figures

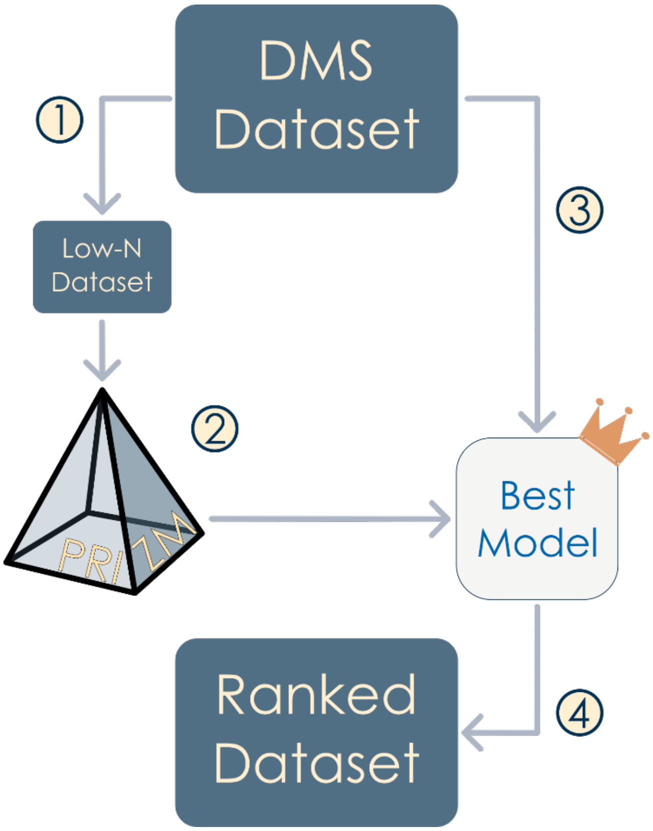


**Figure S1 – Validation approach for PRIZM using low-N datasets from full DMS datasets.** 1) Subsampling of full DMS datasets from ProteinGym (Notin *et al.* 2023) to generate low-N dataset; 2) identification of the best model for low-N dataset; 3) processing of full DMS dataset using best model; 4) ranking of DMS dataset using zero-shot scores and subsequent performance evaluation.

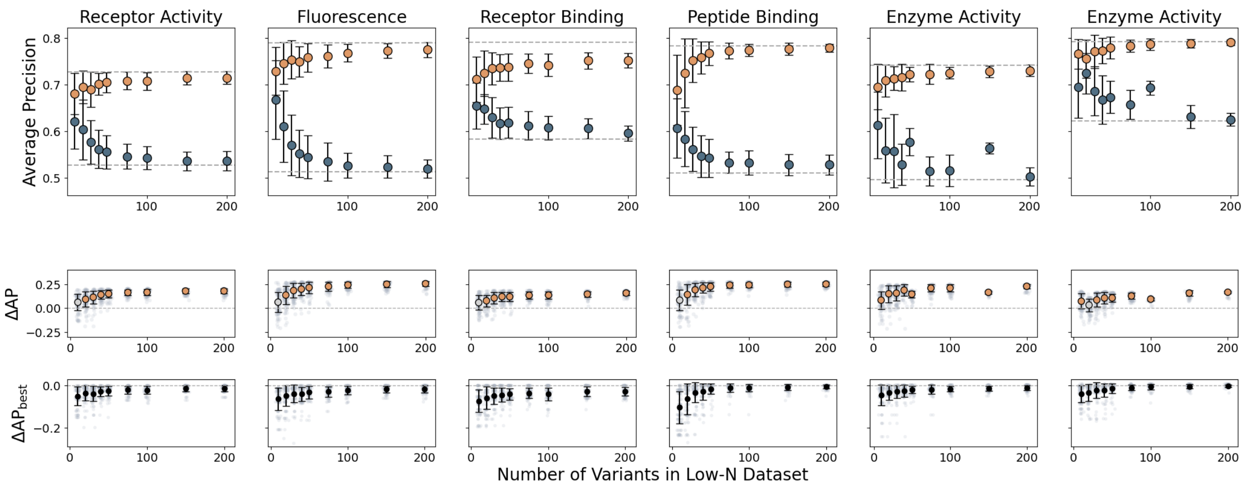


**Figure S2 – Performance of zero-shot models selected by PRIZM as a function of the number of variants in the low-N dataset.** Figure shows performance on full DMS datasets, with fluorescence data from Ellis et al. (Ellis *et al.* 2024), receptor binding from Starr et al. (Starr *et al.* 2020), peptide binding from Araya et al. (Araya *et al.* 2012), and enzyme activity from Romero et al. (Romero, Tran, and Abate 2015) and Chiasson et al. (Chiasson *et al.* 2020). The top row shows average precision (AP) for the best (orange) and worst (blue) models, with dotted lines indicating the overall best and worst models. The middle row shows ΔAP between the best and worst models, colored by Cohen's d (blue: $d < 0.2$, grey: $d < 0.5$, orange: $d > 0.5$). The bottom row shows ΔAP between each selected model and the best overall model ($\Delta AP_{best}$). Error bars represent standard deviation from 100 PRIZM iterations.

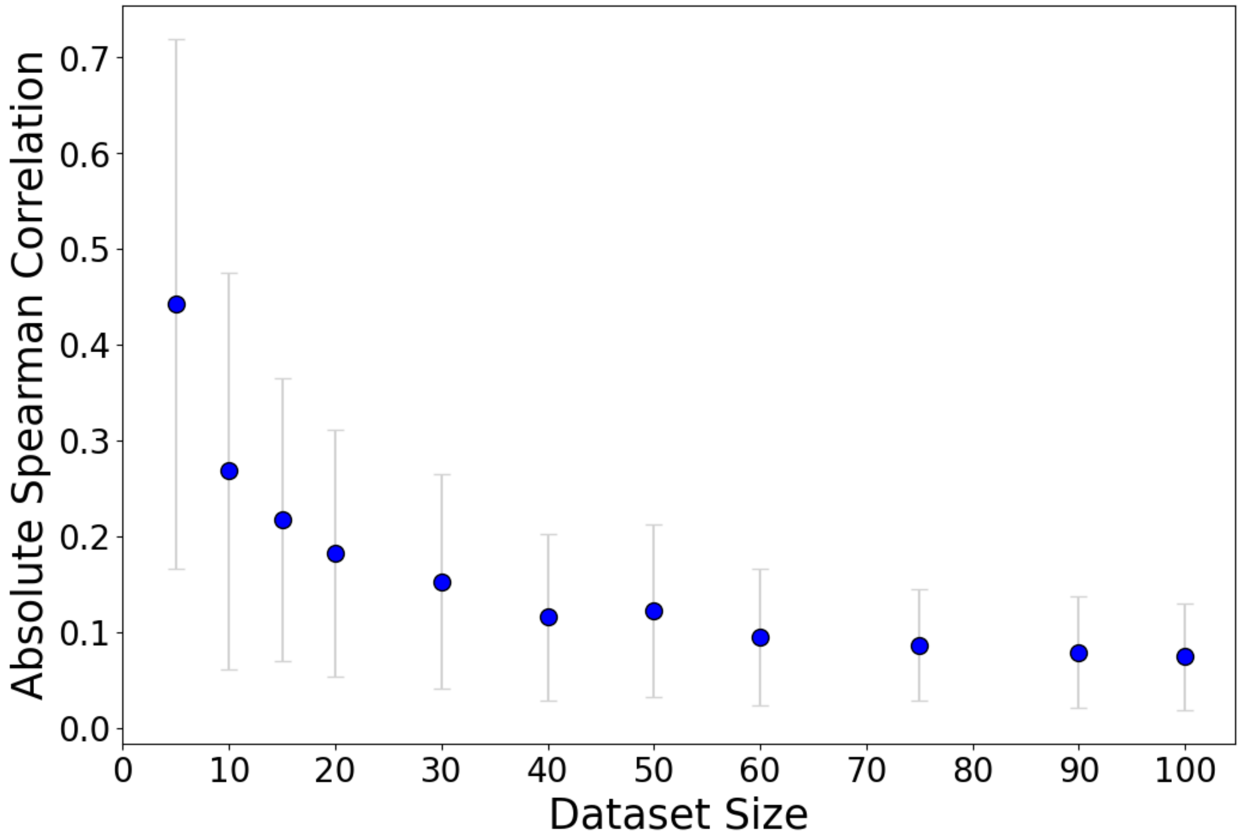


**Figure S3 – Spearman correlation as a function of dataset size.** Average absolute Spearman correlation of randomly generated datasets as a function of their sizes. The average is based on 100 datasets, while the error bars indicate the estimated standard deviation.

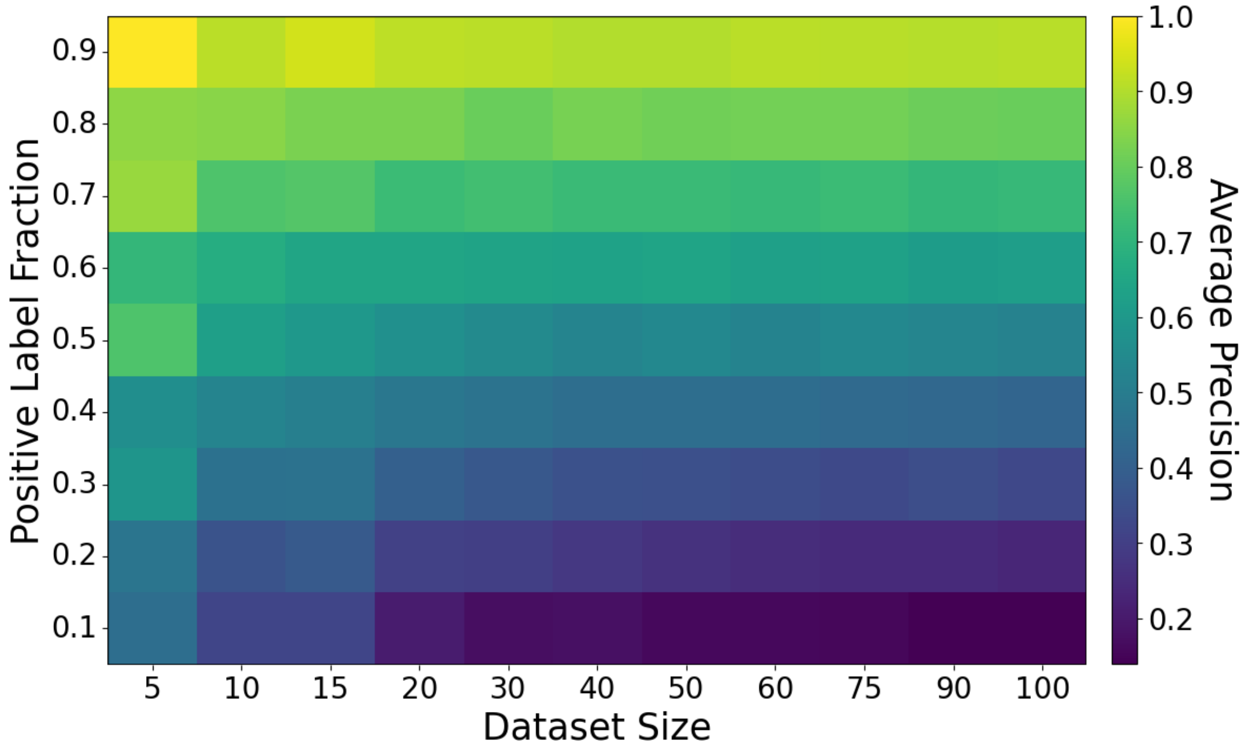


**Figure S4 – Average precision as a function of dataset size and positive label fraction.** Mean average precision across randomly generated datasets as a function of their sizes and positive label fraction. The mean is based on 100 datasets, while the error bars indicate the estimated standard deviation.

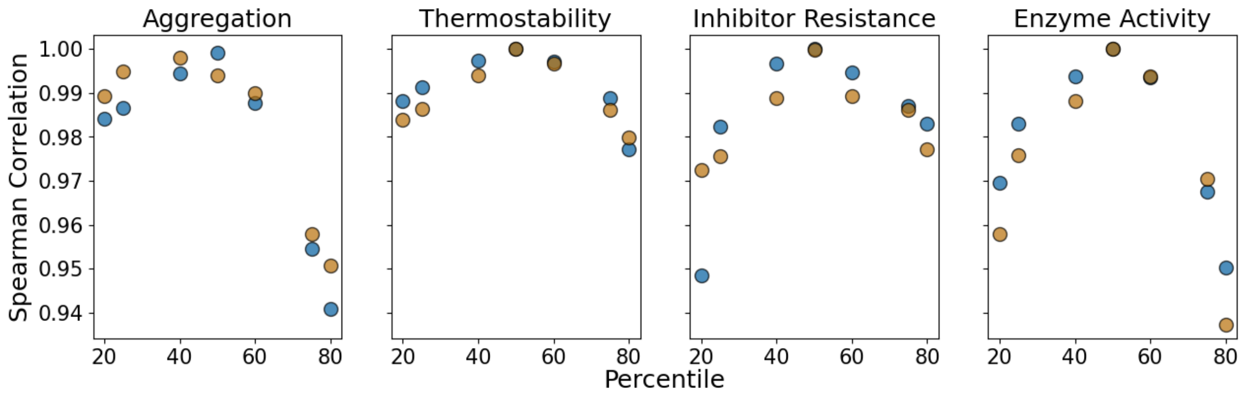


**Figure S5 — Sensitivity of model ranking to binarization threshold.** Spearman correlation between model rankings obtained using percentile-based thresholds and rankings obtained using the default threshold from ProteinGym (Notin *et al*. 2023), shown for four representative benchmark datasets (Brenan *et al*. 2016, Nutschel *et al*. 2020, Hobbs *et al*. 2022, Seuma, Lehner, and Bolognesi 2022). Blue points represent model rankings for the full dataset, while orange points denote average rankings for 100 iterations of subsampling of 50 variants.

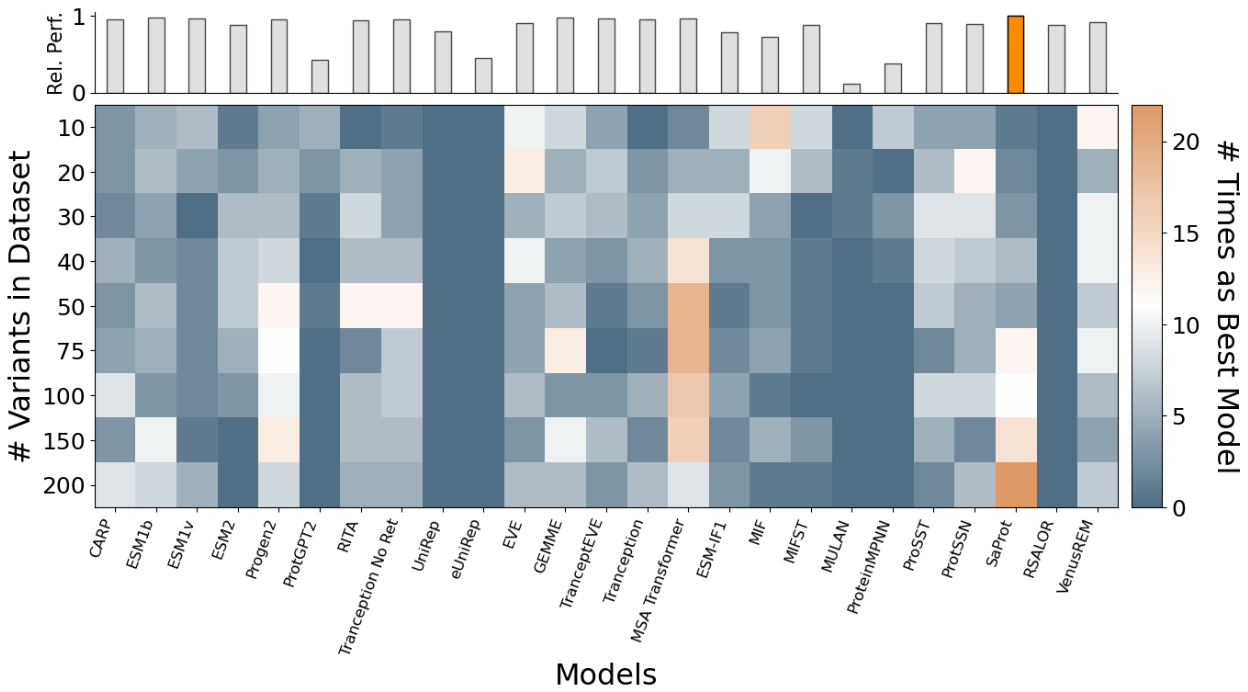


**Figure S6 – Frequency of each model being selected as the best by PRIZM for receptor activity from Jones et al.** (Jones *et al.* 2020)**.** The figure shows data from 100 iterations of the PRIZM workflow. The top inset shows each model's average precision relative to the overall best model (orange) on the full dataset.

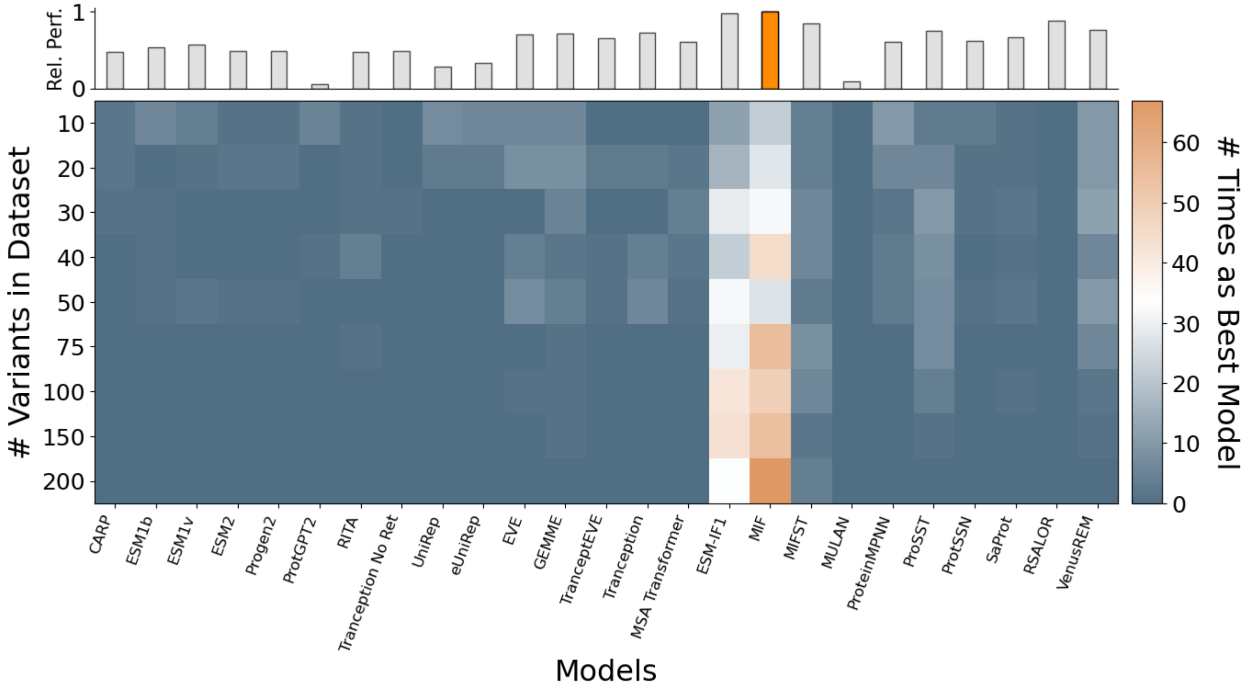


**Figure S7 – Frequency of each model being selected as the best by PRIZM for thermostability from Nutschel et al.** (Nutschel *et al.* 2020)**.** The figure shows data from 100 iterations of the PRIZM workflow. The top inset shows each model's average precision relative to the overall best model (orange) on the full dataset.

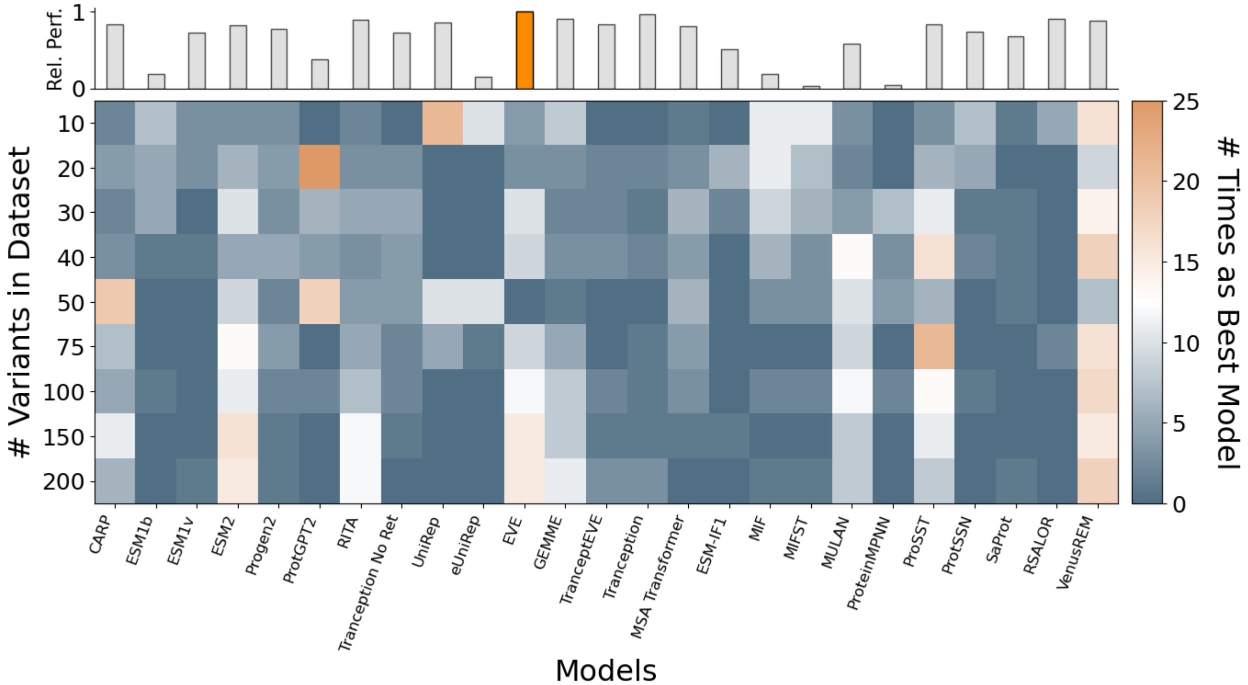


**Figure S8 – Frequency of each model being selected as the best by PRIZM for inhibitor resistance from Brenan et al.** (Brenan *et al*. 2016)**.** The figure shows data from 100 iterations of the PRIZM workflow. The top inset shows each model's average precision relative to the overall best model (orange) on the full dataset.

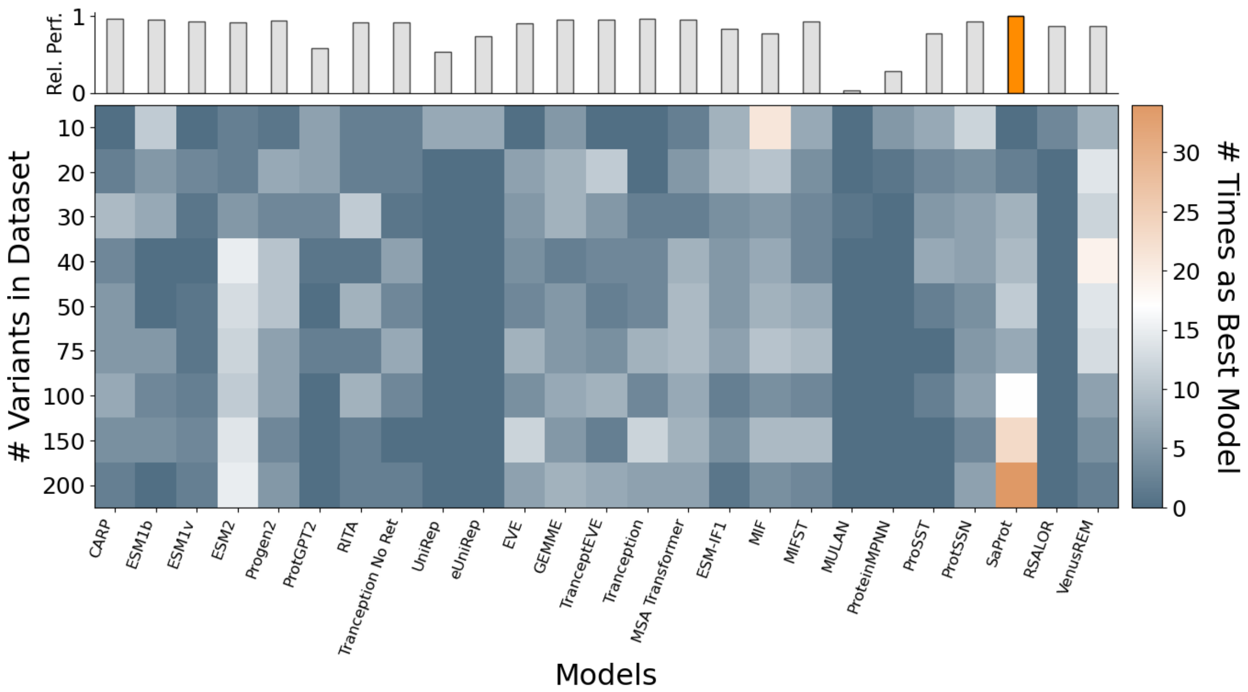


**Figure S9 – Frequency of each model being selected as the best by PRIZM for fluorescence from Ellis et al.** (Ellis *et al.* 2024)**.** The figure shows data from 100 iterations of the PRIZM workflow. The top inset shows each model's average precision relative to the overall best model (orange) on the full dataset.

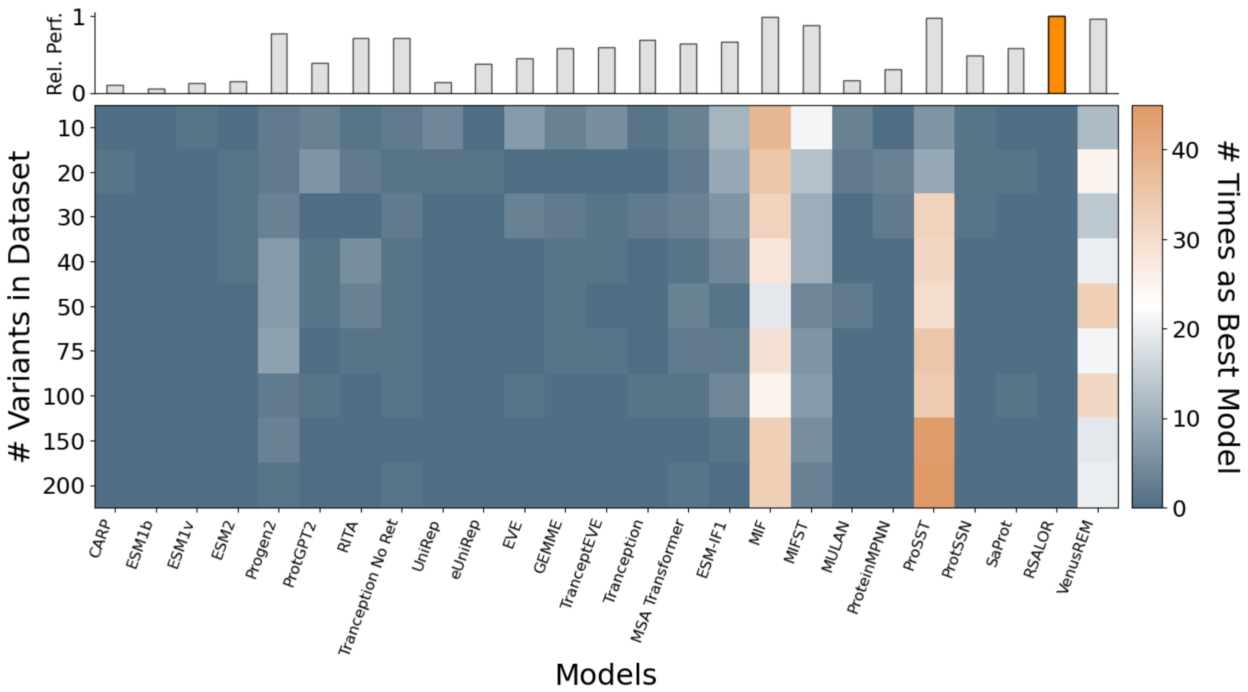


**Figure S10 – Frequency of each model being selected as the best by PRIZM for receptor binding from Starr et al.** (Starr *et al.* 2020)**.** The figure shows data from 100 iterations of the PRIZM workflow. The top inset shows each model's average precision relative to the overall best model (orange) on the full dataset.

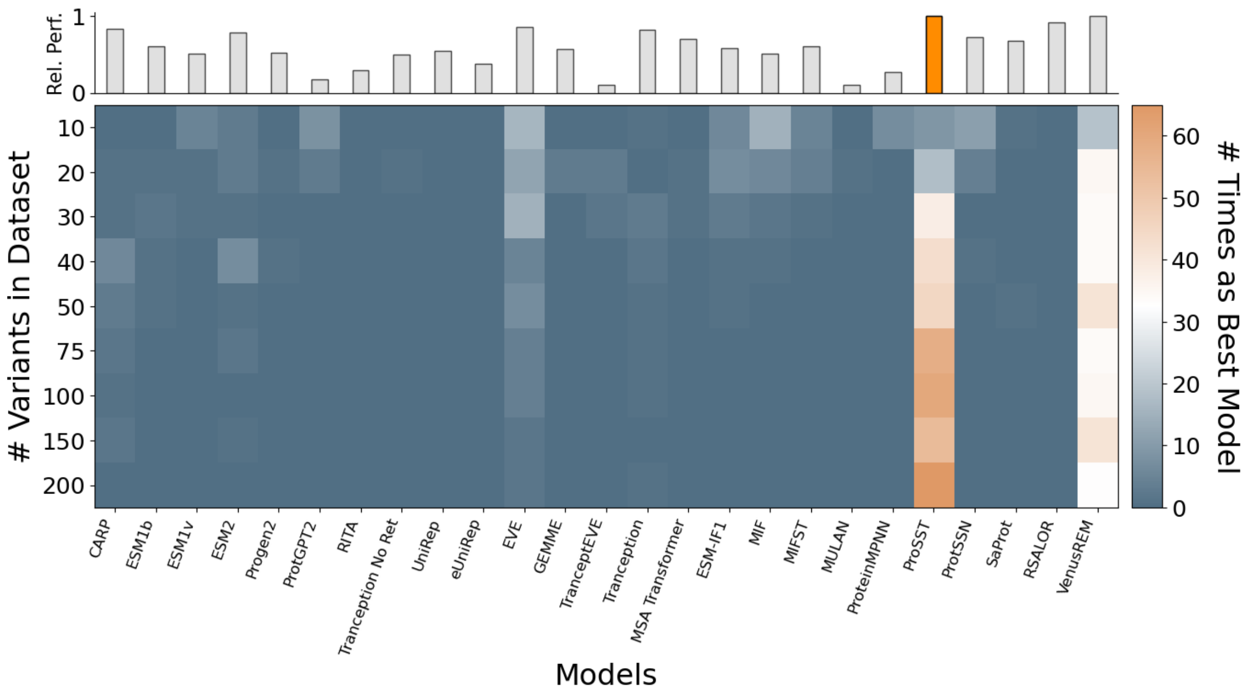


**Figure S11 – Frequency of each model being selected as the best by PRIZM for peptide binding from Araya et al.** (Araya *et al.* 2012)**.** The figure shows data from 100 iterations of the PRIZM workflow. The top inset shows each model's average precision relative to the overall best model (orange) on the full dataset.

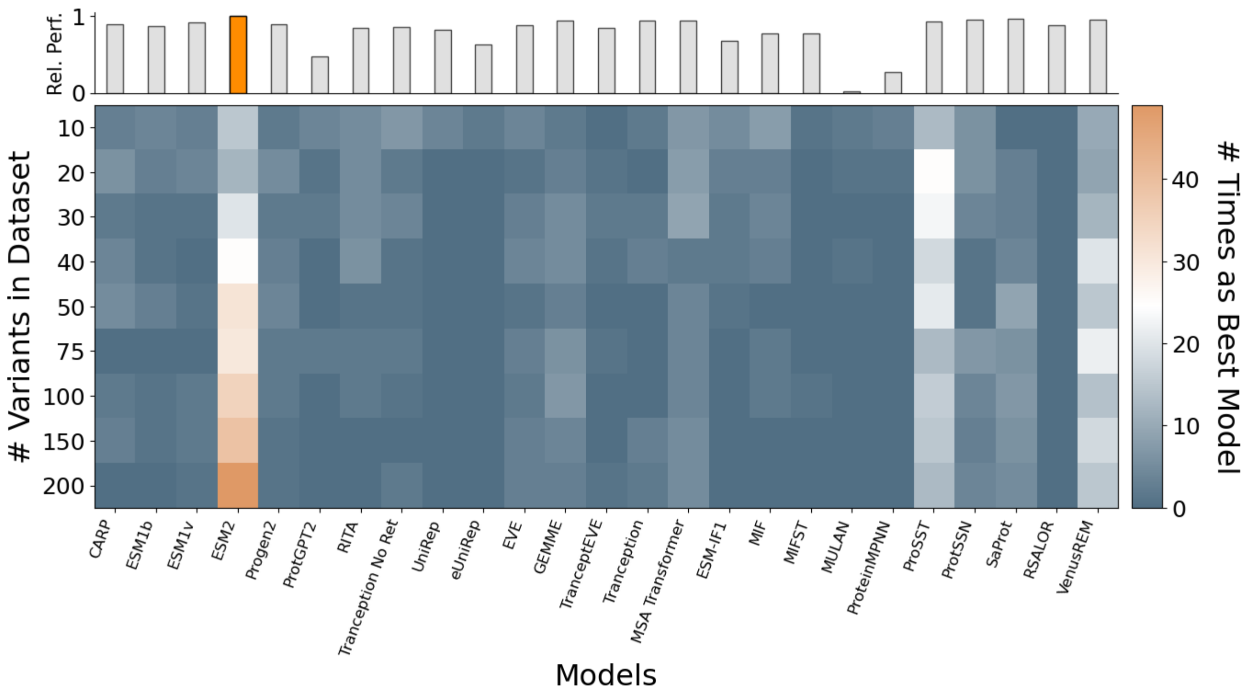


**Figure S12 – Frequency of each model being selected as the best by PRIZM for peptide binding from Hobbs et al.** (Hobbs *et al.* 2022)**.** The figure shows data from 100 iterations of the PRIZM workflow. The top inset shows each model's average precision relative to the overall best model (orange) on the full dataset.

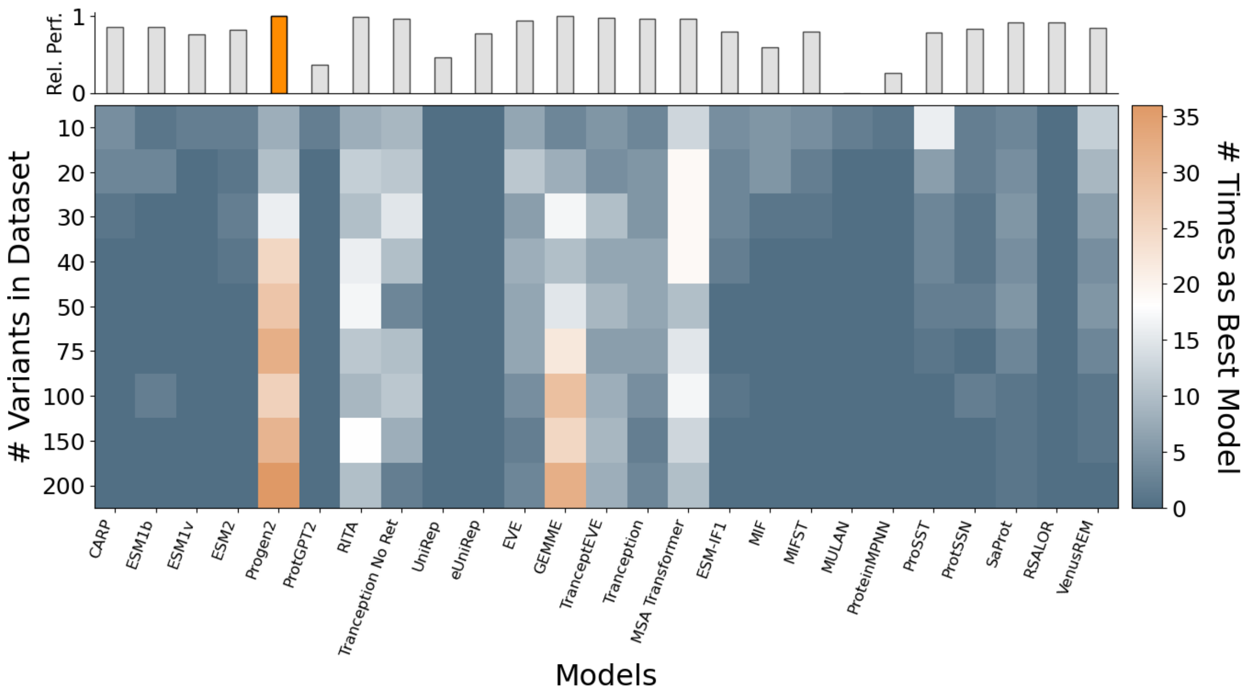


**Figure S13 – Frequency of each model being selected as the best by PRIZM for peptide binding from Romero et al.** (Romero, Tran, and Abate 2015)**.** The figure shows data from 100 iterations of the PRIZM workflow. The top inset shows each model's average precision relative to the overall best model (orange) on the full dataset.

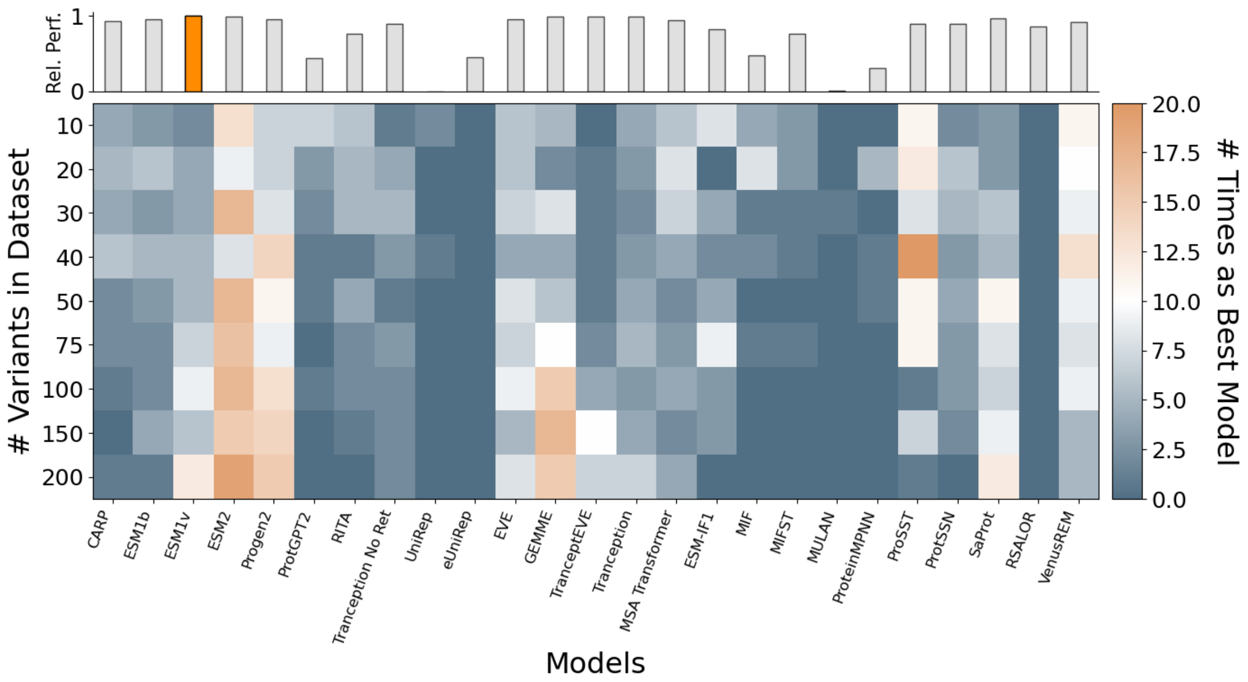


**Figure S14 – Frequency of each model being selected as the best by PRIZM for peptide binding from Chiasson et al.** (Chiasson *et al*. 2020)**.** The figure shows data from 100 iterations of the PRIZM workflow. The top inset shows each model's average precision relative to the overall best model (orange) on the full dataset.

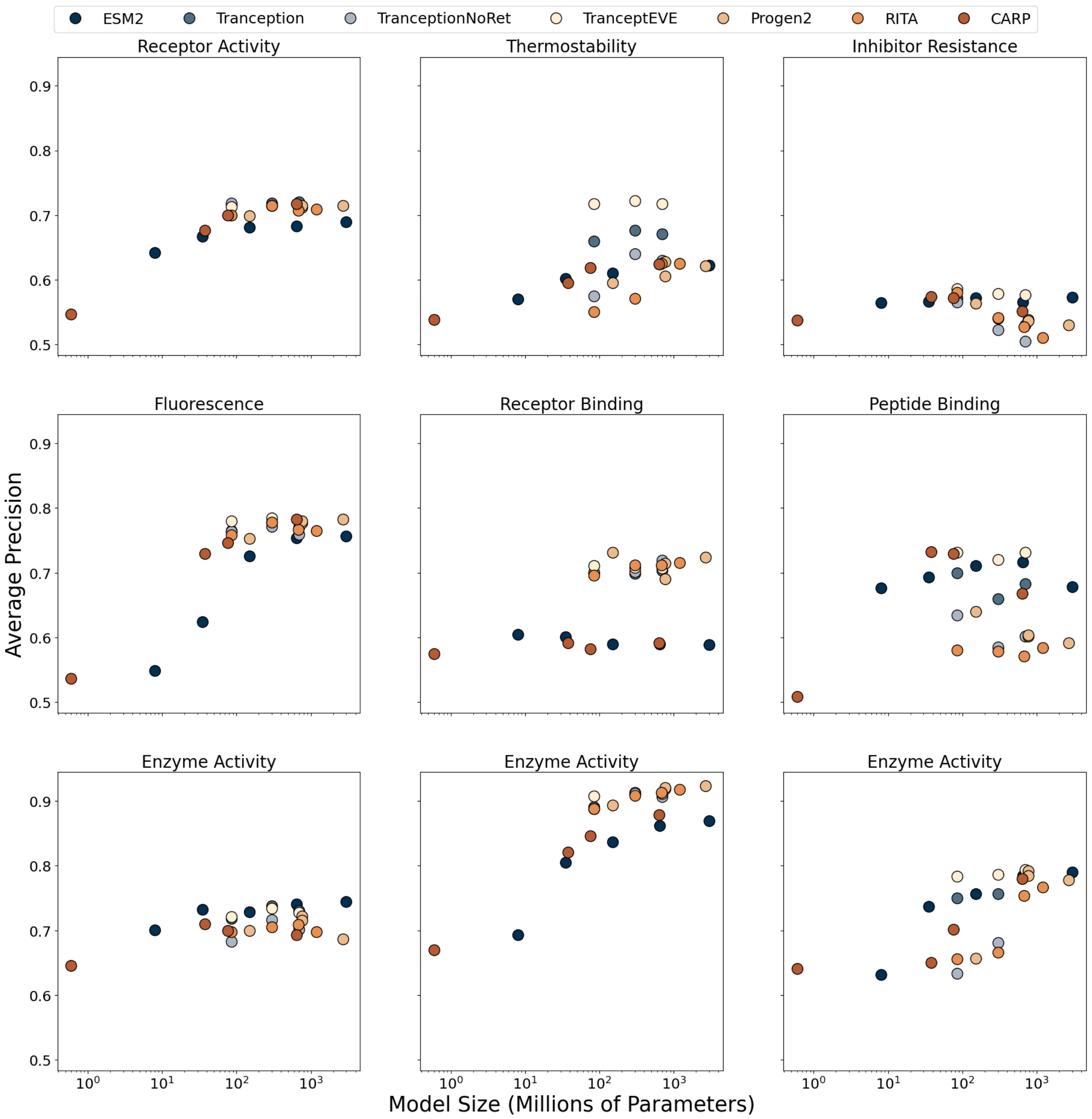


**Figure S15 – Average precision of models as a function of model size.** Figure shows performance on full DMS datasets, with receptor activity data from Jones et al. (Jones *et al.* 2020), thermostability from Nutschel et al. (Nutschel *et al.* 2020), inhibitor resistance from Brenan et al. (Brenan *et al.* 2016) , fluorescence data from Ellis et al. (Ellis *et al.* 2024), receptor binding from Starr et al. (Starr *et al.* 2020), peptide binding from Araya et al. (Araya *et al.* 2012), and enzyme activity from Hobbs et al. (Hobbs *et al.* 2022), Romero et al. (Romero, Tran, and Abate 2015), and Chiasson et al. (Chiasson *et al.* 2020).

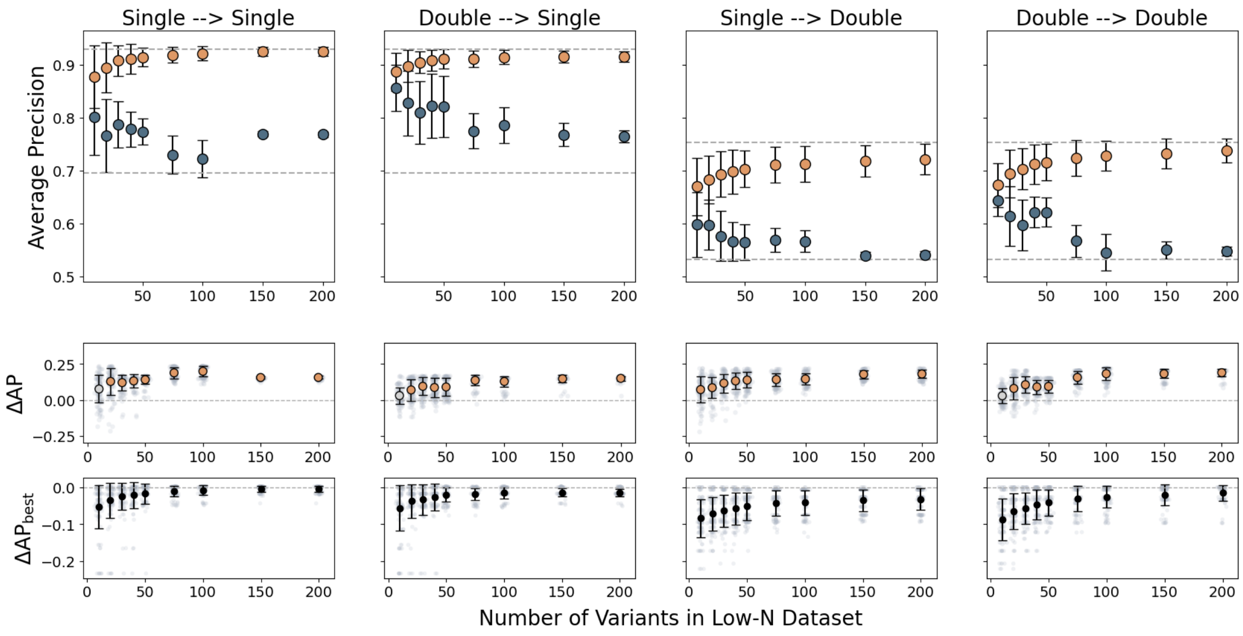


**Figure S16 – Performance of zero-shot models selected by PRIZM using single or double mutants for predicting aggregation data** (Seuma, Lehner, and Bolognesi 2022)**.** "Single → Double" indicates Model Selection using only single mutants and Variant Ranking evaluation on double mutants; other combinations follow the same scheme. The top row shows average precision (AP) for the best (orange) and worst (blue) models, with dotted lines indicating the overall best and worst models. The middle row shows ΔAP between the best and worst models, colored by Cohen's d (blue: $d < 0.2$, grey: $d < 0.5$, orange: $d > 0.5$). The bottom row shows ΔAP between each selected model and the best overall model ($\Delta AP_{best}$). Error bars represent standard deviation from 100 PRIZM iterations.

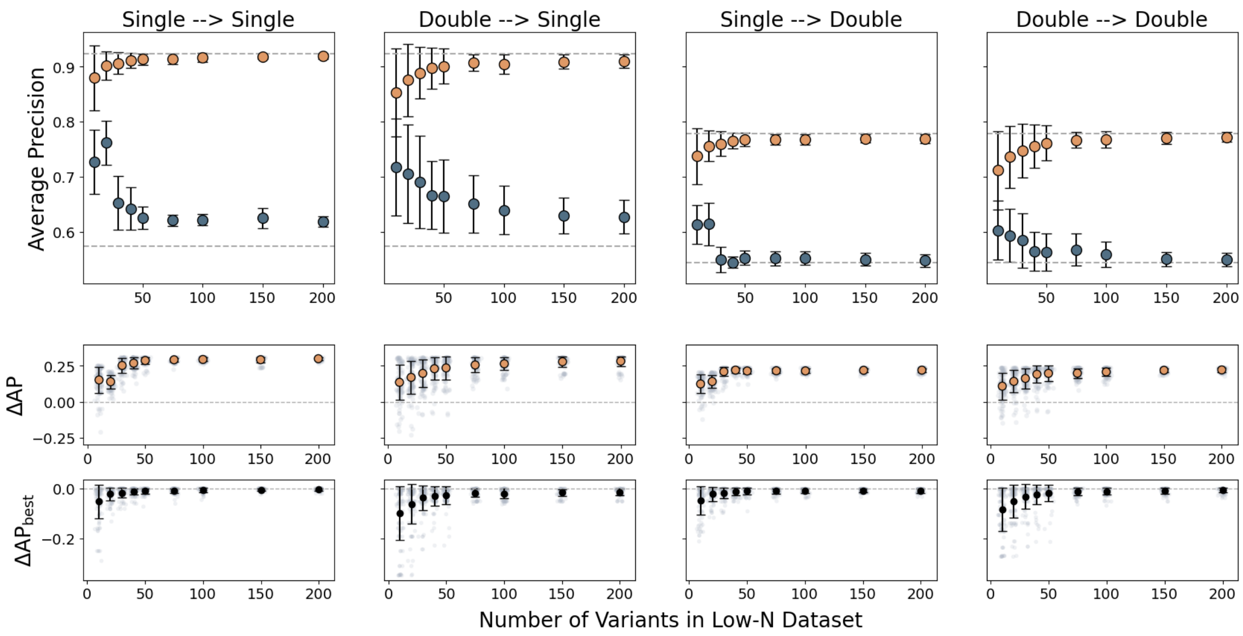


**Figure S17 – Performance of zero-shot models selected by PRIZM using single or double mutants for predicting peptide binding data** (Araya *et al.* 2012). "Single → Double" indicates Model Selection using only single mutants and Variant Ranking evaluation on double mutants; other combinations follow the same scheme. The top row shows average precision (AP) for the best (orange) and worst (blue) models, with dotted lines indicating the overall best and worst models. The middle row shows ΔAP between the best and worst models, colored by Cohen's d (blue: $d < 0.2$, grey: $d < 0.5$, orange: $d > 0.5$). The bottom row shows ΔAP between each selected model and the best overall model ($\Delta AP_{best}$). Error bars represent standard deviation from 100 PRIZM iterations.

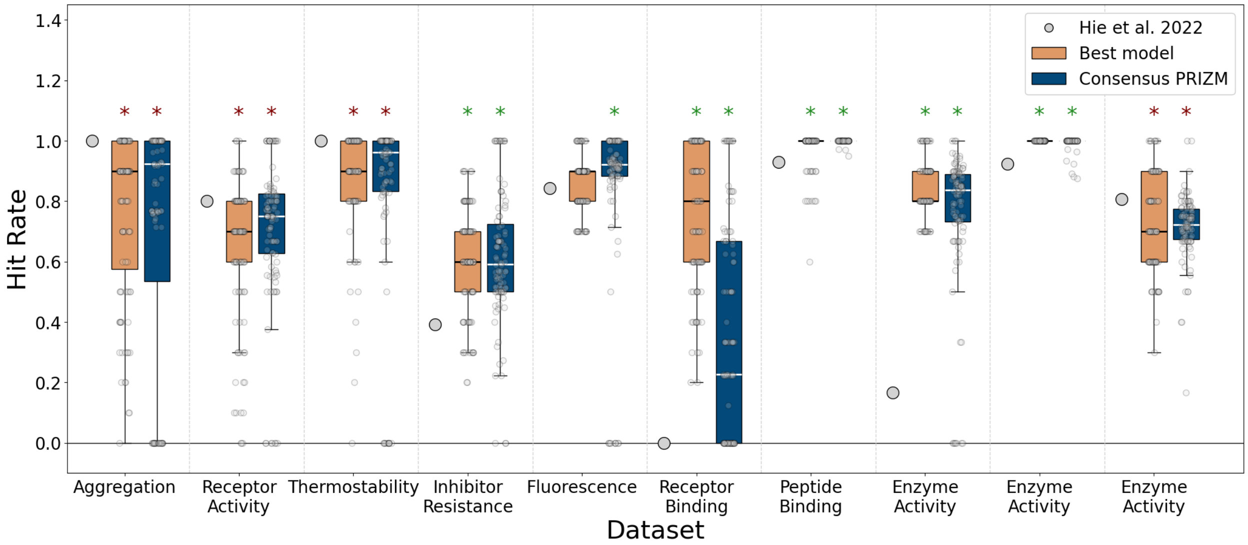


**Figure S18 – Hit rate of best models identified by PRIZM compared to the approach by Hie et al.** (Hie *et al.* 2024)**.** Box plots show the 25th, 50th (median), and 75th percentiles from 100 PRIZM iterations using 50 variants. Gray circles indicate individual hit rates for each method. Asterisks denote significant differences ($p$ = 0.0001) based on a one-sided Mann–Whitney U test (green = PRIZM significantly better than Hie et al.; red = significantly worse).

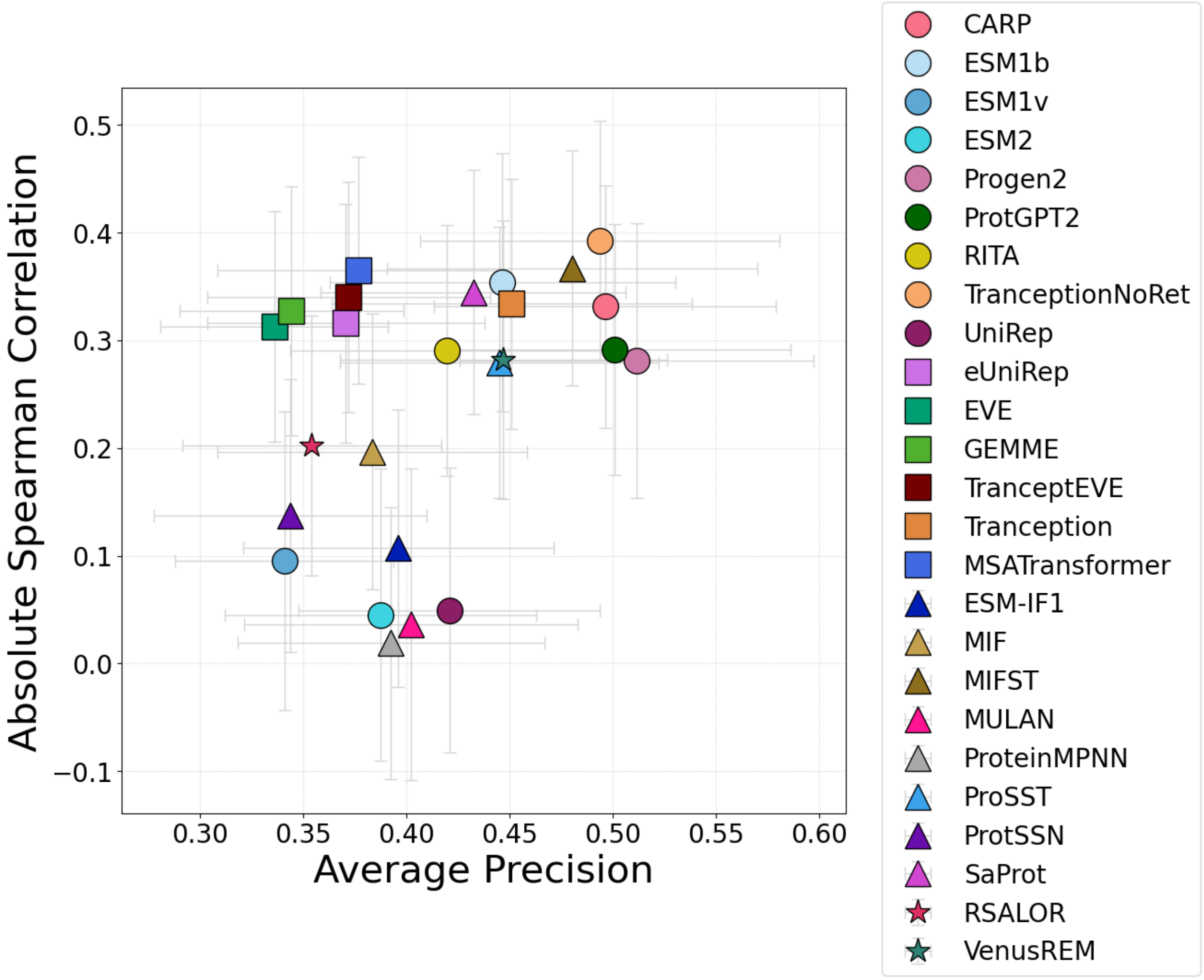


**Figure S19 – Performance metrics for the PRIZM zero-shot models when predicting *Gm*SuSy $T_{m,\ app}$.** Circle = sequence-based model, square = MSA-based model, triangle = structure-based model, star = all three. Error bars indicate the standard deviation using a bootstrapping approach with N = 1000.

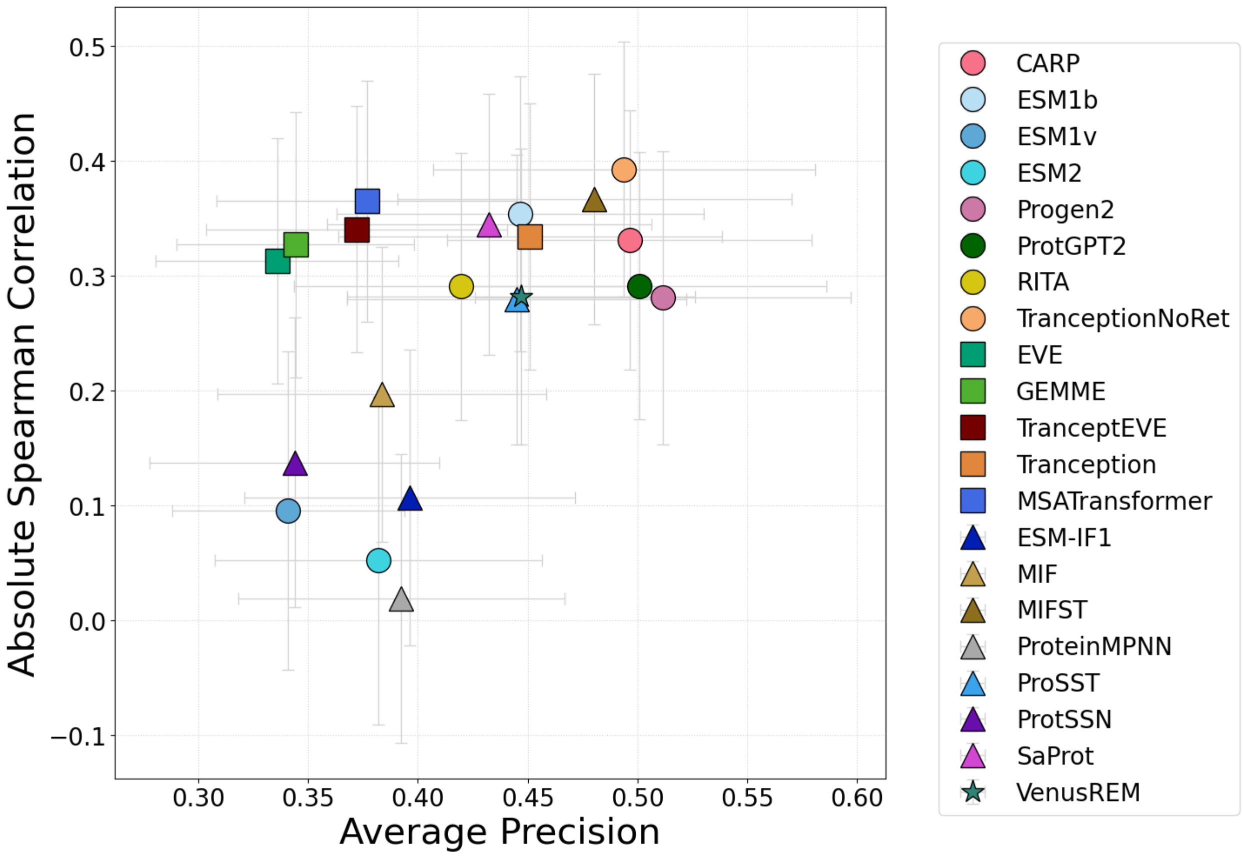


**Figure S20 – Performance metrics for the PRIZM zero-shot models with correct direction when predicting *Gm*SuSy $T_{m,\ app}$.** Only models predicting the correct direction of the mutational effect (e.g., higher values corresponding to improved variants) are shown. Circle = sequence-based model, square = MSA-based model, triangle = structure-based model, star = all three. Error bars indicate the estimated standard deviation using a bootstrapping approach (N = 1000).

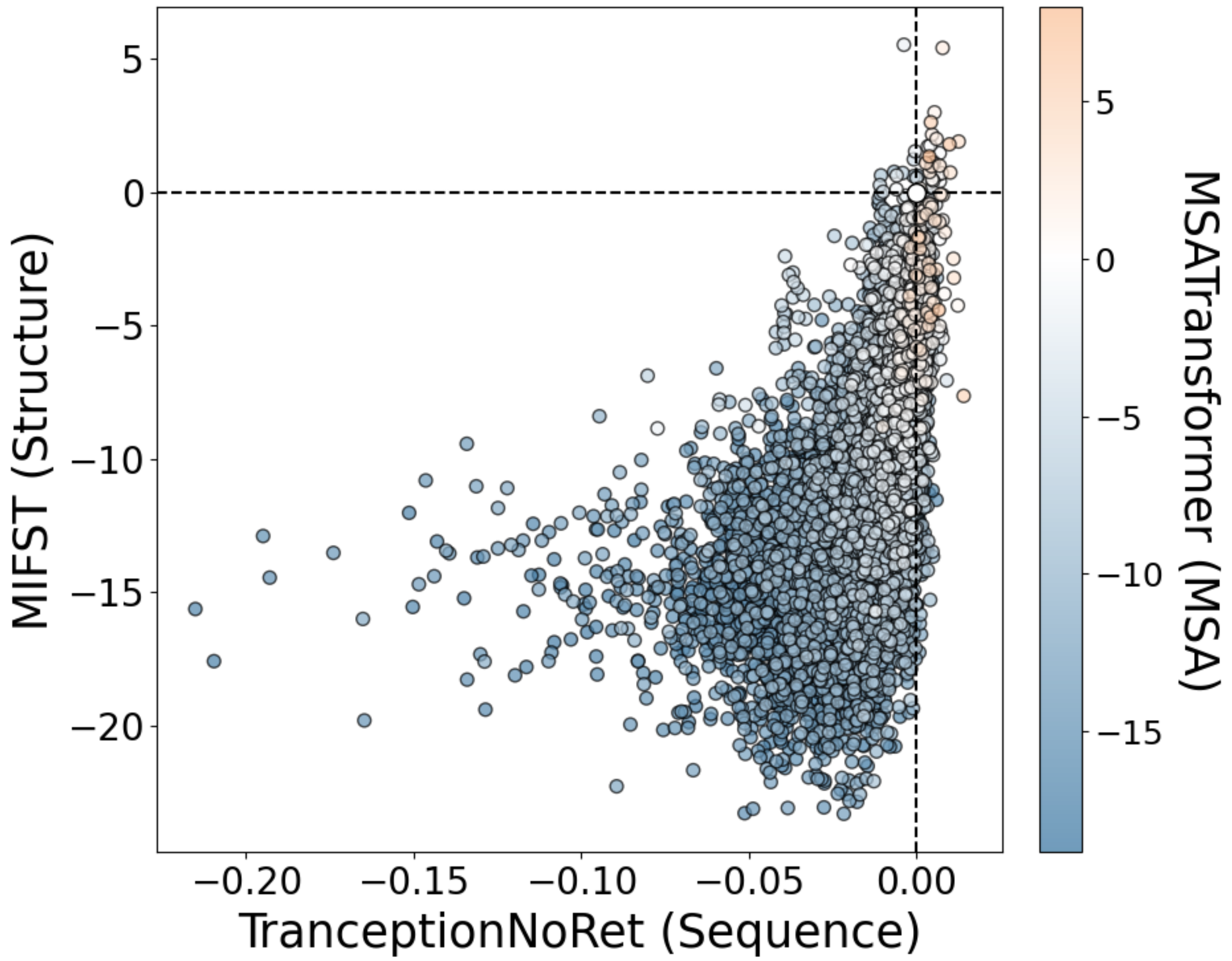


**Figure S21 – Zero-shot scores for *in silico* library of *Gm*SuSy.** Zero-shot scores for best sequence-based model (Tranception No Retrieval) vs. best structure-based model (MIFST) vs. best MSA-based model (MSA Transformer). Dotted lines and white color denote zero-shot scores for WT, while positive values indicate variants better than WT.

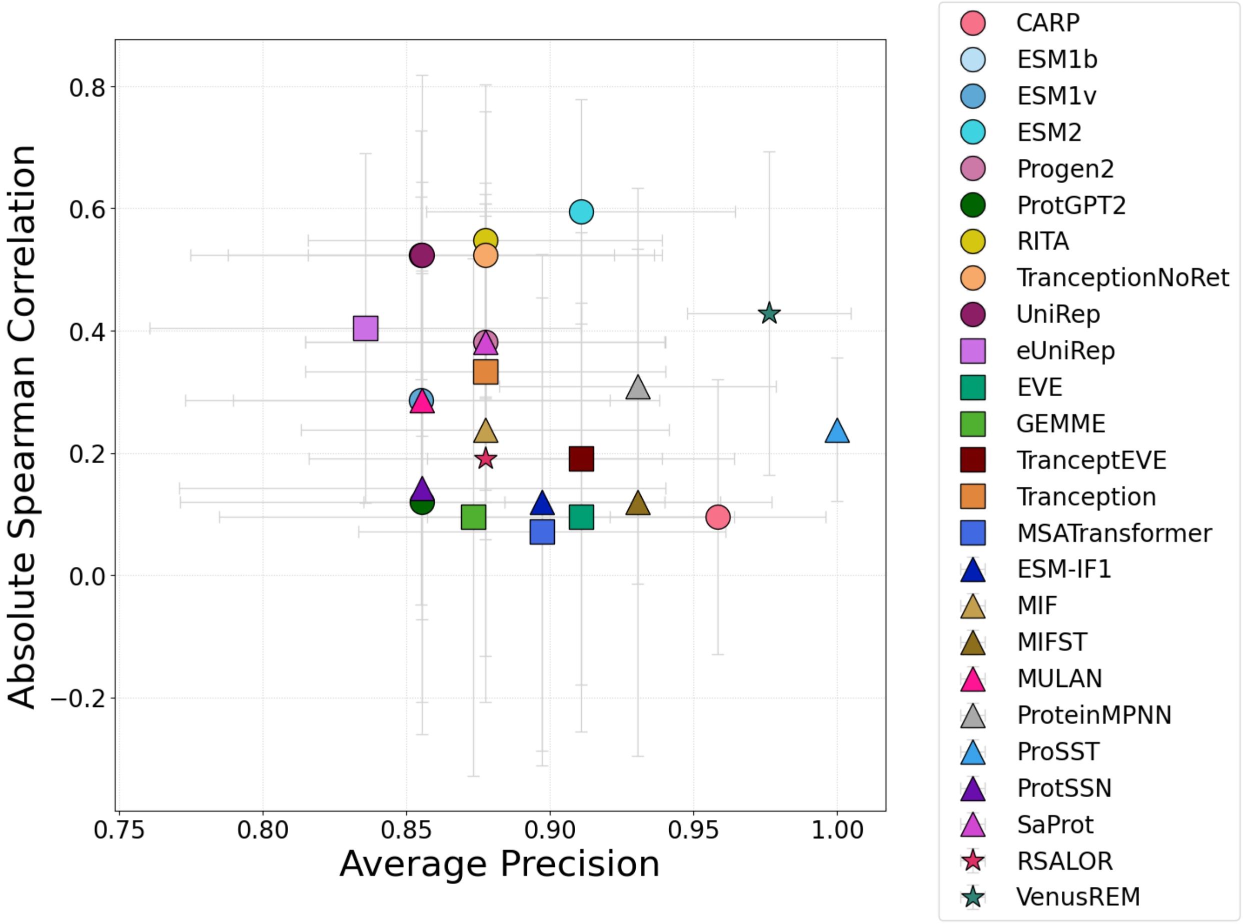


**Figure S22 – Performance metrics for the PRIZM zero-shot models when predicting TOGT1_1 relative activity.** Circle = sequence-based model, square = MSA-based model, triangle = structure-based model, star = all three. Error bars indicate the estimated standard deviation using a bootstrapping approach (N = 1000).

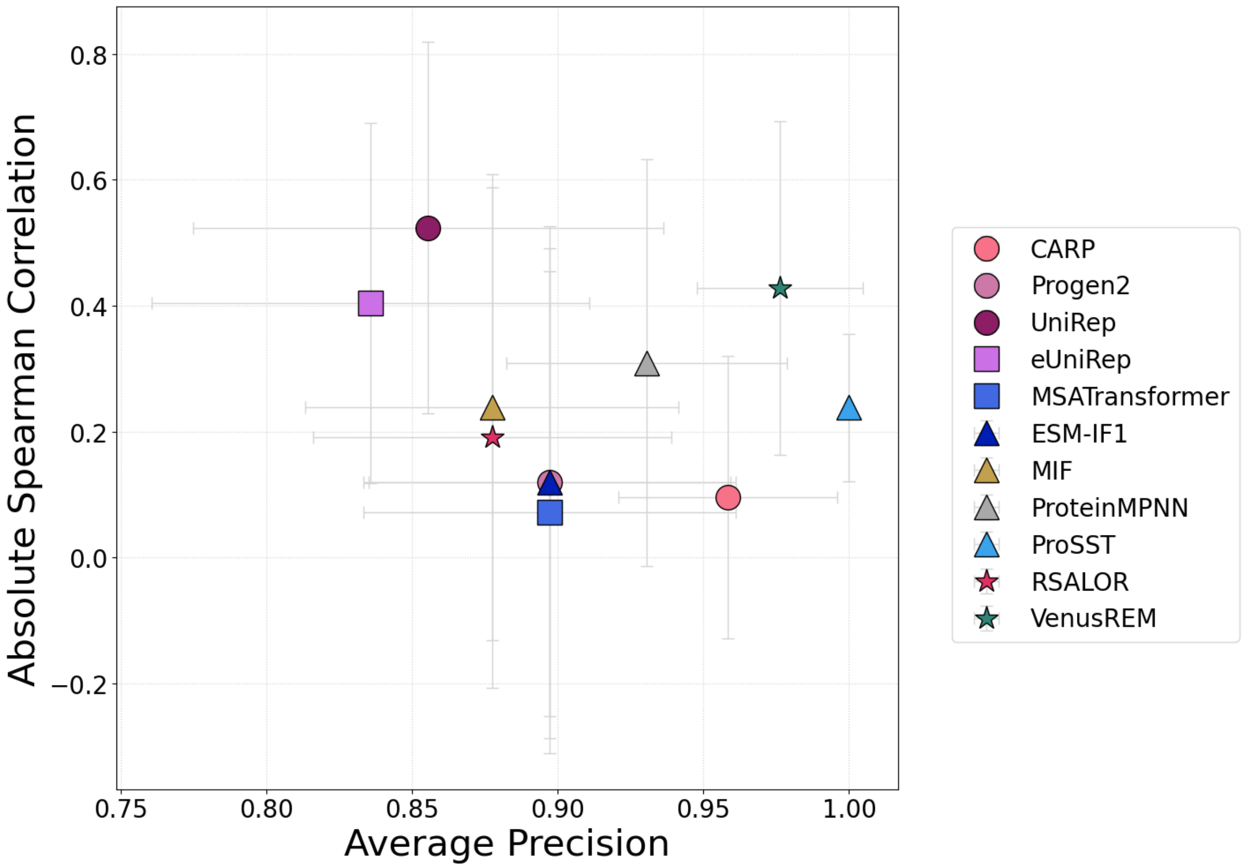


**Figure S23 – Performance metrics for the PRIZM zero-shot models with correct direction when predicting TOGT1_1 relative activity.** Only models predicting the correct direction of the mutational effect (e.g., higher values corresponding to improved variants) are shown. Circle = sequence-based model, square = MSA-based model, triangle = structure-based model, star = all three. Error bars indicate the estimated standard deviation using a bootstrapping approach (N = 1000).

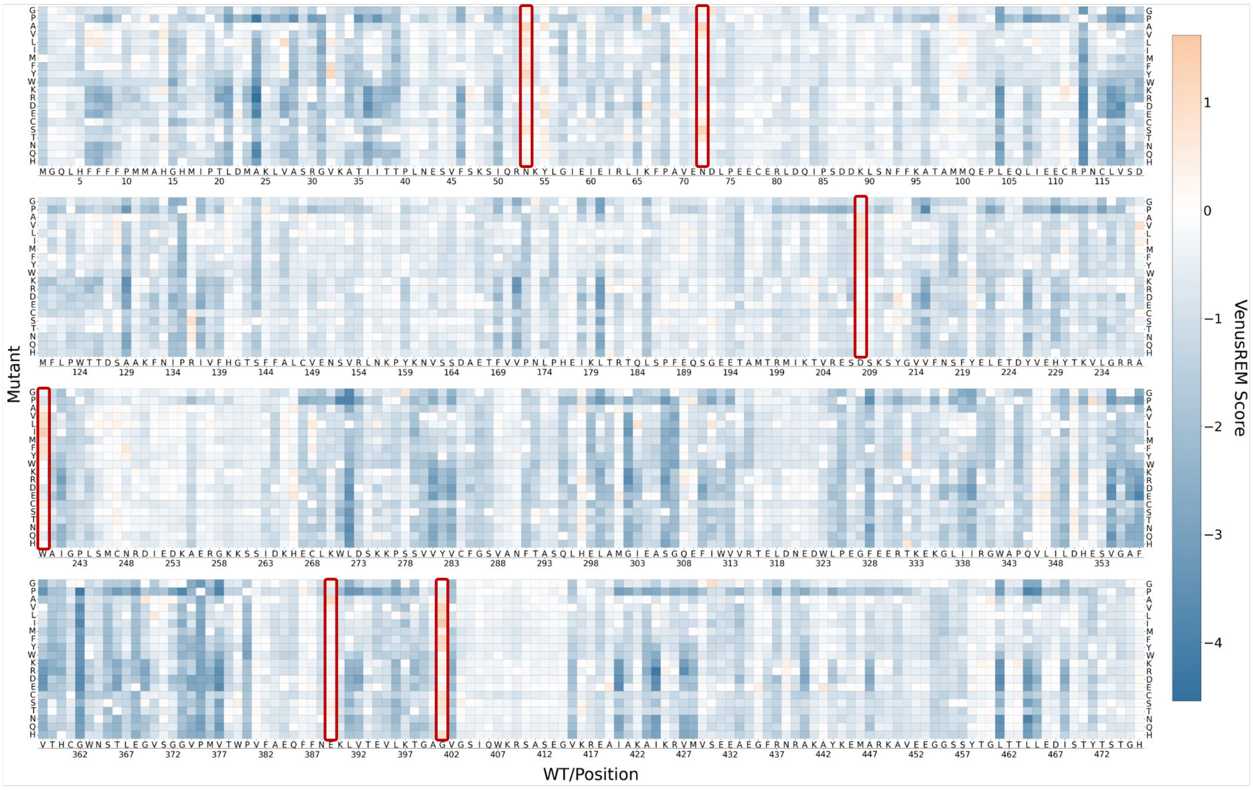


**Figure S24 – TOGT1_1 in silico landscape of VenusREM scores.** Orange denotes variants predicted to be better than WT, while blue denotes variants predicted to be worse. Red squares mark positions selected for experimental characterization due to high proposed mutability.

#### Supplementary Tables

**Table S1 – Zero-shot models contained in the PRIZM collection.**

| Model | Reference | Code[a] |
|---|---|---|
| *Sequence* | | |
| CARP | (Yang, Fusi, and Lu 2024) | https://github.com/microsoft/protein-sequence-models |
| ESM-1b | (Rives *et al.* 2021) | https://github.com/facebookresearch/esm |
| ESM-1v | (Meier *et al.* 2021) | https://github.com/facebookresearch/esm |
| ESM-2 | (Lin *et al.* 2023) | https://github.com/facebookresearch/esm |
| ProGen2 | (Nijkamp *et al.* 2023) | https://github.com/salesforce/progen |
| ProtGPT2 | (Ferruz, Schmidt, and Höcker 2022) | https://huggingface.co/nferruz/ProtGPT2 |
| RITA | (Hesslow *et al.* 2022) | https://github.com/lightonai/RITA |
| Tranception No Retrieval | (Notin, Dias *et al.* 2022) | https://github.com/OATML-Markslab/Tranception |
| UniRep | (Alley *et al.* 2019) | https://github.com/churchlab/UniRep<br>https://github.com/chloechsu/combining-evolutionary-and-assay-labelled-data |
| *MSA* | | |
| eUniRep | (Biswas *et al.* 2021) | https://github.com/churchlab/UniRep<br>https://github.com/chloechsu/combining-evolutionary-and-assay-labelled-data |
| EVE | (Frazer *et al.* 2021) | https://github.com/OATML-Markslab/EVE |
| GEMME | (Laine, Karami, and Carbone 2019) | https://hub.docker.com/r/elodielaine/gemme |
| TranceptEVE | (Notin, Niekerk *et al.* 2022) | https://github.com/OATML-Markslab/ProteinGym |
| Tranception | (Notin, Dias *et al.* 2022) | https://github.com/OATML-Markslab/Tranception |
| MSA Transformer | (Rao *et al.* 2021) | https://github.com/rmrao/msa-transformer |
| *Structure* | | |
| ESM-IF1 | (Hsu *et al.* 2022) | https://github.com/facebookresearch/esm |
| MIF | (Yang, Zanichelli, and Yeh 2023) | https://github.com/microsoft/protein-sequence-models |
| MIFST | (Yang, Zanichelli, and Yeh 2023) | https://github.com/microsoft/protein-sequence-models |
| MULAN | (Frolova *et al.* 2024) | https://github.com/DFrolova/MULAN |
| ProteinMPNN | (Dauparas *et al.* 2022) | https://github.com/dauparas/ProteinMPNN |
| ProSST | (Li *et al.* 2024) | https://github.com/ai4protein/ProSST |
| ProtSSN | (Tan, Zhou *et al.* 2025) | https://github.com/ai4protein/ProtSSN |
| SaProt | (Su *et al.* 2024) | https://github.com/westlake-repl/SaProt |
| *All* | | |
| RSALOR | (Tsishyn *et al.* 2025) | https://github.com/3BioCompBio/RSALOR |
| VenusREM | (Tan, Wang *et al.* 2025) | https://github.com/ai4protein/VenusREM |

[a] While the "Code" column refers to individual model repositories, all code was leveraged from https://github.com/OATML-Markslab/ProteinGym (Notin *et al.* 2023).

**Table S2 – DMS benchmark datasets used for validating PRIZM.**

| **ProteinGym** (Notin *et al.* 2023) **ID** | **Protein** | **Property** | **Reference** |
|---|---|---|---|
| A4_HUMAN_Seuma_2022 | Amyloid beta | Aggregation | (Seuma, Lehner, and Bolognesi 2022) |
| ADRB2_HUMAN_Jones_2020 | $\beta_2$-adrenergic receptor | Receptor activity (Transcription) | (Jones *et al.* 2020) |
| ESTA_BACSU_Nutschel_2020 | Lipase A | Thermostability | (Nutschel *et al.* 2020) |
| MK01_HUMAN_Brenan_2016 | Mitogen-activated protein kinase 1 | Inhibitor resistance | (Brenan *et al.* 2016) |
| SC6A4_HUMAN_Young_2021 | Sodium-dependent serotonin transporter | Fluorescence | (Ellis *et al.* 2024) |
| SPIKE_SARS2_Starr_2020_binding | SARS-CoV-2 spike receptor binding domain | Receptor binding | (Starr *et al.* 2020) |
| YAP1_HUMAN_Araya_2012 | Human Yes-associated protein | Peptide binding | (Araya *et al.* 2012) |
| ANCSZ_Hobbs_2022 | Tyrosine kinase | Enzyme activity | (Hobbs *et al.* 2022) |
| Q59976_STRSQ_Romero_2015 | β-glucosidase | Enzyme activity | (Romero, Tran, and Abate 2015) |
| VKOR1_HUMAN_Chiasson_2020_activity | Epoxide reductase | Enzyme activity | (Chiasson *et al.* 2020) |

**Table S3 – Experimental $T_{m,app}$ and residual activity for best *Gm*SuSy variants as predicted by PRIZM.**

| Variant | PRIZM Rank[a] | $\Delta T_{m,app}$ | Residual Activity |
|---|---|---|---|
| WT | N/A | 0 ± 0.18 | 23.4 ± 11.1 |
| Q528M | 1 | −0.12 ± 0.17 | 16.7 ± 4.3 |
| V658Y | 2 | 1.45 ± 0.84 | 3.0 ± 4.3 |
| H534Y | 3 | −0.21 ± 0.06 | 26.8 ± 14.1 |
| L731E | 4 | 0.29 ± 0.38 | 21.1 ± 12.6 |
| F468I | 5 | 2.96 ± 0.20 | 60.1 ± 19.5 |

[a] Sequence-wide rank.

**Table S4 – Experimental relative activity for selected TOGT1_1 variants.**

| Variant | PRIZM Rank[a] | Relative Activity |
|---|---|---|
| WT | N/A | 100 ± 4.28 |
| N53A | 2 | 60.0 ± 4.58 |
| N72S | 1 | 110.5 ± 2.22 |
| W239I | 1 | 102.69 ± 4.71 |
| G401F | 3 | 119.9 ± 2.84 |
| G401I | 1 | 114.1 ± 2.03 |

[a] Position-wide rank.

Harding-Larsen D, Madsen CD, Teze D *et al.* GASP: A Pan-Specific Predictor of Family 1 Glycosyltransferase Acceptor Specificity Enabled by a Pipeline for Substrate Feature

Generation and Large-Scale Experimental Screening. *ACS Omega* published online 25 Jun. 2024. https://doi.org/10.1021/ACSOMEGA.4C01583.

Hesslow D, Zanichelli NO, Notin P *et al. RITA: A Study on Scaling Up Generative Protein Sequence Models*. 11 May 2022. https://arxiv.org/pdf/2205.05789 (23 Jun. 2025, date last accessed).

Hie BL, Shanker VR, Xu D *et al.* Efficient evolution of human antibodies from general protein language models. *Nat Biotechnol* 2024;**42**(2):275–83. https://doi.org/10.1038/S41587-023-01763-2.

Hobbs HT, Shah NH, Shoemaker SR *et al.* Saturation mutagenesis of a predicted ancestral Syk-family kinase. *Protein Science* 2022;**31**(10). https://doi.org/10.1002/PRO.4411.

Hsu C, Verkuil R, Liu J *et al.* Learning inverse folding from millions of predicted structures. *BioRxiv* 6 Sep. 2022:2022.04.10.487779. https://doi.org/10.1101/2022.04.10.487779.

Jiang K, Yan Z, Bernardo M Di *et al.* Rapid in silico directed evolution by a protein language model with EVOLVEpro. *Science (1979)* 2025;**387**(6732). https://doi.org/10.1126/SCIENCE.ADR6006.

Jones EM, Lubock NB, Venkatakrishnan AJ *et al.* Structural and functional characterization of G protein-coupled receptors with deep mutational scanning. *Elife* 2020;**9**:1–52. https://doi.org/10.7554/ELIFE.54895.

Laine E, Karami Y, Carbone A. GEMME: A Simple and Fast Global Epistatic Model Predicting Mutational Effects. *Mol Biol Evol* 2019;**36**(11):2604–19. https://doi.org/10.1093/MOLBEV/MSZ179.

Lax BM, García ME, Fernandez-Antunez C *et al.* Enzymatic Glucosylation Enhances the Solubility of Niclosamide but Abrogates Its Therapeutic Efficacy. *ACS Omega* published online 26 Jan. 2026. https://doi.org/10.1021/ACSOMEGA.5C12185.

Li M, Tan Y, Ma X *et al.* ProSST: Protein Language Modeling with Quantized Structure and Disentangled Attention. *Adv Neural Inf Process Syst* 2024;**37**:35700–26. https://github.com/ai4protein/ProSST (20 Oct. 2025, date last accessed).

Lin Z, Akin H, Rao R *et al.* Evolutionary-scale prediction of atomic-level protein structure with a language model. *Science (1979)* 2023;**379**(6637):1123–30. https://doi.org/10.1126/SCIENCE.ADE2574.

Meier J, Rao R, Verkuil R *et al.* Language models enable zero-shot prediction of the effects of mutations on protein function. *Adv Neural Inf Process Syst* 2021;**34**:29287–303. https://github. (23 Jun. 2025, date last accessed).

Mejia-Otalvaro F, Delima D, Mendonça C *et al.* Stability engineering of sucrose synthase for robust UDP-glucose regeneration. *BioRxiv* 31 Jan. 2026:2026.01.30.702896. https://doi.org/10.64898/2026.01.30.702896.

Nijkamp E, Ruffolo JA, Weinstein EN *et al.* ProGen2: Exploring the boundaries of protein language models. *Cell Syst* 2023;**14**(11):968-978.e3. https://doi.org/10.1016/j.cels.2023.10.002.

Notin P, Dias M, Frazer J *et al.* Tranception: Protein Fitness Prediction with Autoregressive Transformers and Inference-time Retrieval. Preprint, PMLR, 28 Jun. 2022, 16990–7017. https://proceedings.mlr.press/v162/notin22a.html (23 Jun. 2025, date last accessed).

Notin P, Kollasch A, Ritter D *et al.* ProteinGym: Large-Scale Benchmarks for Protein Fitness Prediction and Design. *Adv Neural Inf Process Syst* 2023;**36**:64331–79.

Notin P, Niekerk L Van, Kollasch AW *et al.* TranceptEVE: Combining Family-specific and Family-agnostic Models of Protein Sequences for Improved Fitness Prediction. *BioRxiv* 27 Dec. 2022:2022.12.07.519495. https://doi.org/10.1101/2022.12.07.519495.

Nutschel C, Fulton A, Zimmermann O *et al.* Systematically Scrutinizing the Impact of Substitution Sites on Thermostability and Detergent Tolerance for Bacillus subtilis Lipase A. *J Chem Inf Model* 2020;**60**(3):1568–84. https://doi.org/10.1021/ACS.JCIM.9B00954.

Rao RM, Liu J, Verkuil R *et al.* MSA Transformer. Preprint, PMLR, 1 Jul. 2021, 8844–56. https://proceedings.mlr.press/v139/rao21a.html (20 Oct. 2025, date last accessed).

Rives A, Meier J, Sercu T *et al.* Biological structure and function emerge from scaling unsupervised learning to 250 million protein sequences. *Proceedings of the National Academy of Sciences* 2021;**118**(15):e2016239118. https://doi.org/10.1073/PNAS.2016239118.

Romero PA, Tran TM, Abate AR. Dissecting enzyme function with microfluidic-based deep mutational scanning. *Proc Natl Acad Sci U S A* 2015;**112**(23):7159–64. https://doi.org/10.1073/PNAS.1422285112.

Seuma M, Lehner B, Bolognesi B. An atlas of amyloid aggregation: the impact of substitutions, insertions, deletions and truncations on amyloid beta fibril nucleation. *Nat Commun* 2022;**13**(1):1–13. https://doi.org/10.1038/S41467-022-34742-3.

Starr TN, Greaney AJ, Hilton SK *et al.* Deep Mutational Scanning of SARS-CoV-2 Receptor Binding Domain Reveals Constraints on Folding and ACE2 Binding. *Cell* 2020;**182**(5):1295-1310.e20. https://doi.org/10.1016/j.cell.2020.08.012.

Su J, Han C, Zhou Y *et al.* SaProt: Protein Language Modeling with Structure-aware Vocabulary. *International Conference on Representation Learning* 2024;**2024**:6987–7009. https://github.com/ (20 Oct. 2025, date last accessed).

Tan Y, Wang R, Wu B *et al.* From high-throughput evaluation to wet-lab studies: advancing mutation effect prediction with a retrieval-enhanced model. *Bioinformatics* 2025;**41**(Supplement_1):i401–9. https://doi.org/10.1093/BIOINFORMATICS/BTAF189.

Tan Y, Zhou B, Zheng L *et al.* Semantical and geometrical protein encoding toward enhanced bioactivity and thermostability. *Elife* 2025;**13**. https://doi.org/10.7554/ELIFE.98033.4.

Teufel F, Kollasch AW, Huang Y *et al.* Few-shot Protein Fitness Prediction via In-context Learning and Test-time Training. *ArXiv* 2 Dec. 2025. http://arxiv.org/abs/2512.02315 (31 Mar. 2026, date last accessed).

Tsishyn M, Hermans P, Rooman M *et al.* Residue conservation and solvent accessibility are (almost) all you need for predicting mutational effects in proteins. *Bioinformatics* 2025;**41**(6). https://doi.org/10.1093/BIOINFORMATICS/BTAF322.

Yang KK, Fusi N, Lu AX. Convolutions are competitive with transformers for protein sequence pretraining. *Cell Syst* 2024;**15**(3):286-294.e2. https://doi.org/10.1016/J.CELS.2024.01.008.

Yang KK, Zanichelli N, Yeh H. Masked inverse folding with sequence transfer for protein representation learning. *Protein Eng Des Sel* 2023;**36**. https://doi.org/10.1093/PROTEIN/GZAD015.

Zhou Z, Zhang L, Yu Y *et al.* Enhancing efficiency of protein language models with minimal wet-lab data through few-shot learning. *Nat Commun* 2024;**15**(1):5566. https://doi.org/10.1038/s41467-024-49798-6.
